# Markov Field network model of multi-modal data predicts effects of immune system perturbations on intravenous BCG vaccination in macaques

**DOI:** 10.1101/2024.04.13.589359

**Authors:** Shu Wang, Amy J Myers, Edward B Irvine, Chuangqi Wang, Pauline Maiello, Mark A Rodgers, Jaime Tomko, Kara Kracinovsky, H. Jacob Borish, Michael C Chao, Douaa Mugahid, Patricia A Darrah, Robert A Seder, Mario Roederer, Charles A Scanga, Philana Ling Lin, Galit Alter, Sarah M Fortune, JoAnne L Flynn, Douglas A Lauffenburger

## Abstract

Analysis of multi-modal datasets can identify multi-scale interactions underlying biological systems, but can be beset by spurious connections due to indirect impacts propagating through an unmapped biological network. For example, studies in macaques have shown that BCG vaccination by an intravenous route protects against tuberculosis, correlating with changes across various immune data modes. To eliminate spurious correlations and identify critical immune interactions in a public multi-modal dataset (systems serology, cytokines, cytometry) of vaccinated macaques, we applied Markov Fields (MF), a data-driven approach that explains vaccine efficacy and immune correlations via multivariate network paths, without requiring large numbers of samples (i.e. macaques) relative to multivariate features. Furthermore, we find that integrating multiple data modes with MFs helps to remove spurious connections. Finally, we used the MF to predict outcomes of perturbations at various immune nodes, including a B-cell depletion that induced network-wide shifts without reducing vaccine protection, which we validated experimentally.

## Introduction

In recent decades, various biological fields have explored the benefits of integrating multiple experimental modalities to characterize individual samples from a biological system^1^. Various computational methods have been developed to procure insights into biological phenomena by specifically leveraging the multi-modality of data^2,3^, resulting in improved biomarker discovery, molecular interaction inferences, and predictive power^1^. In principle, the unique advantage of multi- modal data is the ability to measure multiple biological scales, compartments, or aspects that each could play a vital role in the mechanistic or “causal” chains of a biological system, thereby enabling data-driven and relatively less biased identification of the key interactions governing particular phenomena. However, the precise capacity of multi-modal data for distinguishing mechanistic interactions from simple correlations is often not explicitly evaluated, and only implicit in the improved performance of multi- modal analyses.

Many tools can be used for modeling networks of interactions from data, ranging from statistical networks^4^ to spatiotemporal differential equation models^5^. In particular, network modeling can help distinguish when correlations arise from “direct” relations (e.g. interaction) or are an “indirect” consequence (e.g. correlated but not causal) of various pathways within the network (**Figure 1A**). To this end, the general category of methods in Probabilistic Graphical Models (PGMs) (e.g. Bayesian Directed Acyclic Graphs (DAGs) used for causal inference) have the practical benefits of explicitly modeling direct and indirect relations via *conditional dependence*, while requiring relatively modest sample sizes relative to the number of network features, and making minimal mechanistic assumptions. For example, Bayesian DAGs have previously been applied to understand signaling network dynamics^6^ or gene expression^7^, and are intimately related to Structural Equation Modeling approaches used to study disease determinants^8^. Markov Field models, a generalization of Markov Chains and the undirected analogue of Bayesian DAGs, have also been applied to understand ‘omics data in various disease contexts^9^, while also underlying statistical physics and thermodynamics models^10^ of interactions occurring on structured “network” arrangements (e.g. crystalline solids)^11^. A Gaussian Markov Field approach to analyzing metabolomics seemed to learn network interactions enriched for actual metabolic reactions^12^, and a Markov Field was used to briefly describe an integrative analysis of proteomic and metabolomic data of body weight changes in human populations^13^. The main benefit of PGMs is not only in their ability to distinguish direct vs indirect interactions, but also that they model how simultaneous sequences of interactions ultimately lead to outcomes, and so PGMs can help us understand global network effects like compensatory or redundant structures, or how to manipulate parts of a sequence to achieve optimal downstream outcomes. However, while tools for fitting PGMs to datasets are relatively mature, it is often difficult to conceptually and computationally interpret how specific paths, structure, and global topology of a PGM give rise to the statistical correlations between distant features in a biological network.

**Figure 1.**
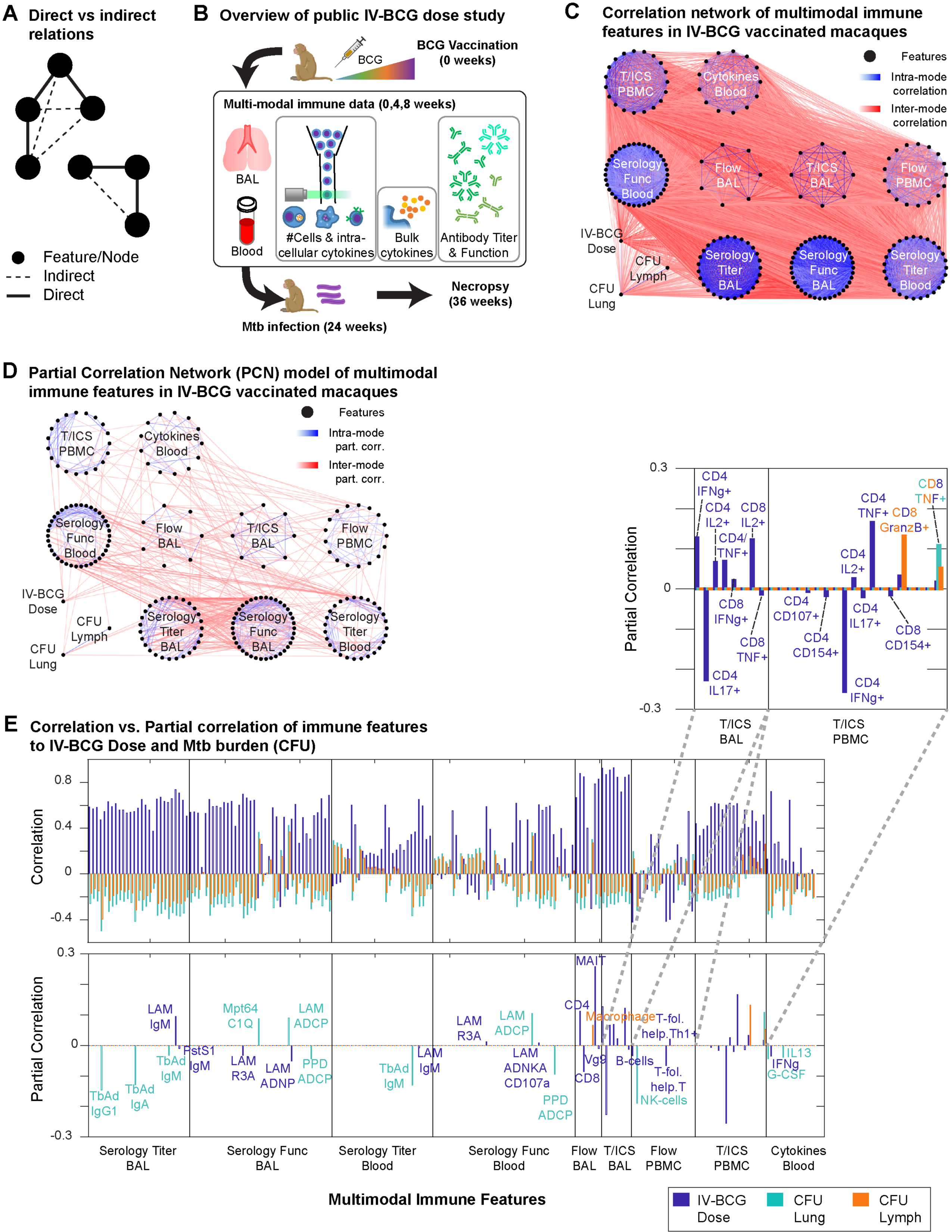
Partial correlation network models of IV-BCG vaccinated macaque immune systems. **A)** Schematic of how “indirect” relations arise from “direct” relations. **B)** Overview of the existing IV-BCG variable dose study, showing the multi-modal features selected for use in our study. **C)** Network of correlations between immune features with magnitude >0.1, with nodes organized by data modes. **D)** Network of partial correlations between immune features fit by Graphical LASSO, for edges with partial correlations of magnitude >0.1. **E)** Correlations and partial correlations of each immune feature with either IV-BCG dose, lung CFU, or lymph node CFU.

**BOX 1 : Progress and potential**

High throughput technologies are now routinely used to generate dense and complex multi- modal datasets across disparate disease and therapeutic spaces. As a consequence, modeling-based approaches are increasingly necessary to discern meaningful patterns in these data and commonly used methods aim to find the features/measurements (e.g., genes in transcriptomic data or antibody titers in serological profiling) that have strongest correlations with a desired outcome. However, these correlates may or may not be the most biologically relevant as the drivers of a desired phenotype, since outcomes are the cumulative response of highly co-correlated features within a larger biological network. Consider an example of a simplified biological network where we have 4 measured features that participate in a vaccine response (Figure). In this system, Cell A (Feature 1) responds to vaccination by becoming activated and secretes Cytokine B (Feature 2) which then stimulates Cell C (Feature 3) to produce high levels of Antibody D (Feature 4); and the antibodies ultimately go on to bind the bacteria and resolve infection. If we did not know the order of our multi-modal features within biological pathways—as is often the case—modeling might identify the strongest correlate of protection as Cell A, an indirect driver of outcome, while missing Antibody D as the feature that actually is the most direct effector of protection. Thus, being able to explicitly model the network of correlations between our measured features and construct pathways between our multi-scale measurements is important for contextualizing whether statistically defined correlations likely represent direct or indirect contributors to outcomes.

In this paper, we empirically dissected the network of correlations between immune features across a multimodal dataset derived from non-human primates that were vaccinated against the bacterial pathogen that causes tuberculosis. This modeling utilized a probabilistic graphical modeling (PGM) approach, Markov Fields using the Graphical Lasso algorithm, in which varying vaccine dose is used as an input and bacterial burden after infection is the outcome, with the multi-modal immune measurements obtained from the animals (comprised of cellular states measured by flow cytometry, cytokine secretion measured by ELISA and antibody titers measured by serological profiling) populating the nodes in-between. By linking immune measurements that have ‘direct’ correlations to each other, we produced a map of how immune features relate to all other datapoints; and discovered several key benefits by using a PGM approach:

1. The model identified frequent connections between features derived from different data modes, indicating applicability of PGM approaches for understanding the relationships between orthogonally generated datapoints across parallel technologies.
2. A majority of measurements within each data type were highly correlated with one another. In contrast, including multi-modal data helped remove confounding predictors without sacrificing resolution, demonstrating that it is preferable to measure multimodal data features across orthogonal technologies than simply including more measurements within a single data type.
3. The preponderance of immune features measured were predicted to be only indirectly connected to vaccination or protection, via relatively few critical paths ultimately linking vaccination with bacterial burden. Interestingly, we also found subsets of immune features directly correlated with protection but unconnected to vaccination, suggesting that a network approach can identify associations both within and external to the specific study intervention being tested. Hence, both vaccine-dependent and -independent correlates of protection conceivably represent targets for vaccine design.
4. Finally, and perhaps most valuably, PGMs offer an opportunity for prospective hypothesis generation by predicting how perturbations of specific immune responses would propagate through the network and impact desired outcomes. In this work, we obtained immunological data from a new vaccination study where animals were also depleted for B cells; and showed that our network model could successfully predict changes in vaccine response that were actually observed in these test animals. This then allowed us to computationally disrupt all the immune response measurements in our network (a scale not experimentally feasible) to identify which immune system changes are expected to abrogate and augment vaccine protection. Such prospective modeling will help translate multimodal data into coherent signals for follow-up validation experiments and hone decision making for future vaccination efforts.

Taken all together, this work exhibits unusual promise in leveraging multi-modal data to answer outstanding mechanistic questions across a wide range of human health and disease states. We have shown that including network modeling through a generalizable PGM framework provides benefits by removing confounding predictors, uncovering novel links between features across technologies and scales, and identifying direct correlates of response. The model also offers us a glimpse of the future, modeling multitudes of potential interventions and identifying the most attractive translational targets for rationally curating the outcomes we want.

To investigate the utility of multi-modal data for identifying key interactions in biological networks, we applied concepts from Markov Fields to a multi-modal dataset of various immune features in macaques vaccinated against tuberculosis with intravenous (IV) BCG^14^, and applied various global graphical analysis techniques to understand how effects of the vaccine propagate through the network of immune features to eventually establish protection against tuberculosis (i.e. correlation between vaccine dose and protection). BCG delivered by the standard intradermal route has limited effectiveness against *Mycobacterium tuberculosis* (Mtb) infection and tuberculosis disease in humans and in macaques^15^.

However, changing the route of BCG administration to IV vaccination resulted in robust protection against infection in disease^16^. Several potential correlates of immunity have been proposed for BCG IV, including T cells and anti-mycobacterial antibodies^17,18^. In a previous study, we undertook a correlates analysis using data from a study in which macaques were vaccinated with a wide range of BCG IV doses, resulting in approximately half the animals showing protection. This study identified CD4 T cells producing cytokines, number of natural killer (NK) cells, and IgA in airways as correlates of protection. These features also had strong associations with multiple other cell types and functions, suggesting a network of immune responses important in BCG IV-induced protection. These and other immune mechanisms could each contribute in unique ways to vaccine efficacy, all be intermediate steps in pathways leading to efficacy, or be downstream “off-target” effects of vaccination that do not contribute to efficacy. Multi-modal data analysis with Markov Fields and global graphical analysis can help distinguish these various possibilities to understand vaccine efficacy.

Herein, we fit and interpret *partial correlation networks* (PCNs), a specific kind of Markov Field, on a multi-modal dataset including antibody titers, antibody-dependent functions, cytokines, and immune cell abundances in macaques vaccinated with IV-BCG^14,19^. We chose PCNs over other PGM methods that assume extra structural constraints, so that any learned networks would be fully data-driven and we could avoid introducing modeling assumptions about how multiple data modes connect to one another.

More broadly, current PGM techniques allowed us to accurately analyze direct vs indirect interactions and network paths even under the practical constraint there were several-fold more features of interest than there were available samples/animals. With PCNs, we were able to determine that the vast majority of correlations between immune features to each other and to disease outcome were indirect and could be explained as the propagated effects along relatively few paths in a sparse multi-modal PCN model. Notably, paths identified by the PCN regularly jump between data modes, reflecting the multi-scale nature of the interactions mediating vaccine efficacy that is unveiled by multi-modal data. Thus, the multi-modal PCN acts as a descriptive tool that can filter out many empirical correlations as potential interactions, based on the definition of conditional dependence. We also applied various global graphical analysis techniques to help interpret the role of different immune correlates to disease outcome, based on their relative position along paths in the PCN. Finally, we show that multi-modal PCNs can be used to systematically model how perturbations to specific features propagate to changes across the whole network. As a validated prediction, we perform an experimental B cell depletion during IV-BCG vaccination in macaques, which induced subtle changes to disease and immune state that could be predicted with reasonable accuracy by the multi-modal PCN. Overall, our work supports the use of Markov Fields and global graphical analysis to distinguish key interactions in multi-modal biological data, and as predictive tools for understanding how perturbations propagate through biological networks.

## Results

### Sparse partial correlation network explains correlations between IV-BCG macaque immune features

The IV-BCG variable dose study contained a cohort of ∼30 rhesus macaques, each measured for an array of humoral and cellular immune features before or after BCG vaccination^14,19^. Whereas previous analyses of the variable dose study focused on describing correlates and predictors of disease outcome in each respective data mode, we aimed to leverage the multi-modal datasets from that same study to identify the immune subnetwork of “direct” interactions (**Figure 1A**) that mediate vaccine-induced effects and disease outcome, in order to achieve predictive understanding of perturbation outcomes. To this end, we chose to analyze 194 immune features measured during the initial weeks post-vaccination (see Methods, **Figure 1B**), chosen based on minimizing the number of missing entries, in conjunction with 3 clinical variables: IV-BCG dose, lung Mtb colony-forming units (lung CFU) and lymph node CFU (LN CFU) measured at time of necropsy. Features represented measurements from bronchoalveolar lavage (BAL)) or blood (e.g. peripheral blood mononuclear cells, PBMCs) compartments, and covered multiple scales of the immune system: antibody titers (Serology Titer), antibody-dependent functions (Serology Func), flow cytometry measurements of cell-type abundances (Flow) and T-cell intra-cellular cytokine expression (T/ICS), and bulk cytokine blood levels (Blood Cytokines). The features were densely correlated, visualized as a correlation network (Fisher transformed Z-test, Storey’s FDR = 0.2, |ρ|>0.1) in **Figure 1C** and **S1A**.

To filter out “indirect” correlations, we instead determined which pairs of features had substantial *partial correlation*, which can be interpreted as “direct” correlation based on the inability of all other features to fully explain the correlation between a feature pair. We fit a partial correlation network (PCN) to all features using the Graphical LASSO algorithm^20,21^ to find network topology, using a threshold of 80% selection stability^22^ and *a posterior* covariance estimation^23^ to fit values of partial correlation (see Methods). We treated measurements from the same monkey at different timepoints (pre-vaccine, 4, 8 weeks post-vaccine) as separate samples, leading to two consequences: 1) statistical power for PCN modeling can be improved due to an increased “sample size” of *N*=98 compared to *N*=30 monkeys, although the gain in power will be less than what would be expected from 98 independent monkeys, since different timepoints from the same monkey are correlated to some degree, and 2) we interpret different timepoints simply as generic measurements of the initial state post-vaccination.

The resulting partial correlation matrix (**Figure S1B**) is visualized in **Figure 1D** for edges with partial correlations greater than 0.1 in magnitude, with only 4% of all possible edges passing this threshold, in contrast to the correlation network in **Figure 1C** that contains ∼60% of all possible edges. Consistent with the density of edges visualized in **Figure 1D**, we estimated that ∼58% of the possible correlations between pairs of immune features are expected to be non-zero by Storey’s FDR. However our PCN model found only 9% of edges to be non-zero (i.e. “direct”), based on our stability selection criteria of 80% stability. Furthermore, using only the sparse set of “direct” correlations, the PCN model was able to adequately explain the “indirect” correlations (R = 0.94, **Figure S1CD**) as the combined effects of various paths within the PCN. Thus, the PCN filtered out ∼90% of correlations as indirect while maintaining the ability to explain them, suggesting that analyzing the direct correlations and the paths they form is sufficient to understand how IV-BCG mediated effects propagate through the macaque immune system lead to protection.

### PCN explains protective effects of vaccine using multiple paths through the immune network

Next, we examined how the PCN helped explain the correlation between initial vaccine dose and eventual Mtb burden (CFU; **Figure S1E**), by first examining their respective direct neighbors in the PCN. Whereas most measured immune features are non-zero correlates (estimated by Storey’s Q FDR) of BCG IV dose (79%), lung CFU (66%), and LN CFU (47%), very few features were “direct” (i.e. partial) correlates in the PCN (**Figure 1E**). Furthermore, while each positive correlate of dose was often simultaneously a negative correlate of CFU, the partial correlates of dose or CFU were almost mutually exclusive, suggesting that Mtb protection is not strictly dependent on IV-BCG dose, but more that vaccination elicits a subset of changes that ultimately propagate through the immune network and limit bacterial load. For example, **Figure 1E** suggests that vaccine dose primarily determines the balance of various types of T-cells and their cytokine expression in both airways and blood, while the direct determinants of CFU are primarily the abundance of macrophages and NK cells in blood (NK cells in BAL had been excluded due to too many missing entries for PCN fitting), alongside specific antibody titers and antibody-dependent cell phagocytosis (ADCP) in both blood and BAL. Thus, the PCN explains vaccine-mediated protection against Mtb as starting with altering T-cell abundance and cytokine expression, but how this ultimately affects Mtb CFU can only be understood by globally dissecting how these effects propagate through the multi-scale immune network.

### Multi-modal data integration leads to sparse PCNs by accounting for inter-mode confounding factors

To understand how the PCN structure was broadly organized at the level of data modes, we asked whether using multiple data modes vs single data modes 1) improved predictive power by introducing more predictors, or 2) changed the paths and topology of PCNs, since it is generally understood that PGM topologies can change substantially when certain nodes are unobserved (i.e. Elimination Algorithm for marginal distributions^24^ demonstrated in **Figure 2A**). First, linear regression on dose, lung CFU, and LN CFU using their respective partial correlates as predictors (**Figure 2B)** was able to explain ∼74-91% of each variable’s variance. More generally, linear regression for any feature/node in the PCN, again only using each node’s respective partial correlates as predictors (often less than 30 of the 196 possible nodes), often explained >80% of a node’s variance (red, **Figure 2C**). Thus, the multi-modal PCN, in spite of its sparsity, had substantial predictive power. For comparison, we also trained a separate PCN using only single data modes plus dose, lung CFU, and LN CFU (single-mode PCNs), following the same stability selection procedure and criteria as was done for the multi-modal PCN to avoid overfitting, and then performed linear regression again in the same manner. In general, the explained variance (R^2^) of each feature (black, **Figure 2C)** was only marginally lower than in the multi-modal PCN (an average decrease in R^2^ of ∼0.05). Thus, multi-modal data integration introduced additional predictors to each node (i.e. neighbors in the PCN), but predictive power did not increase substantially.

**Figure 2.**
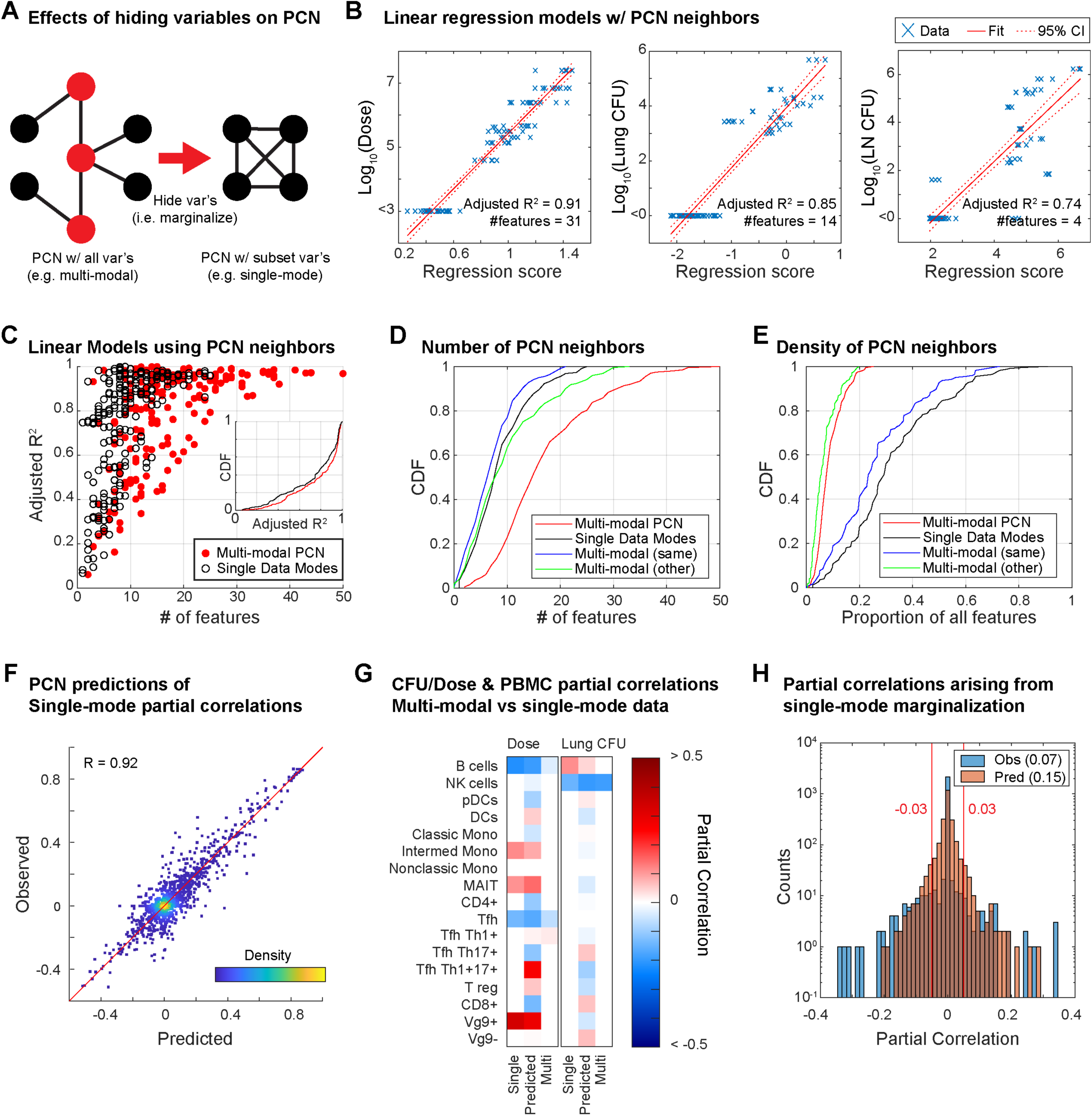
Role of multi-modal data integration in PCN modeling. **A)** Schematic of how partial correlations between nodes can arise if variables are hidden (i.e. the elimination algorithm). Example shows a case in which 4 black nodes with no edges connecting each other become completely connected upon hiding the red nodes. **B)** Linear regression models of dose, lung CFU, and lymph node CFU using only their neighboring features in the PCN. **C)** Performance of linear regression models for predicting any given node using only its neighbors in either the multi-modal PCN or a single-mode PCN. Regression performance is compared with the number of neighboring features. **D)** Cumulative distribution function of number of features neighboring a given node in either multi- or single-mode PCNs, with green and blue showing whether features in the multi-modal PCN arise from the same or different data mode as the node itself. **E)** Same as D, but for the proportion of possible features as opposed to the number. **F)** Observed single-mode partial correlations versus those predicted by the multi-modal PCN. **G)** Partial correlations to dose or lung CFU for the Flow PBMC features from the single-mode PCN, the single-mode PCN predicted by the multi-modal PCN, and the multi-modal PCN. **H)** Distribution of partial correlation values in single-mode PCNs that were previously zero in the multi-modal PCN.

However, multi-modal data did help eliminate spurious connections: although the multi-modal PCN typically had more total edges per feature (red, **Figure 2D**), we found that it had fewer edges within a feature’s corresponding data mode (blue) than the single-mode PCNs (black), indicating that the multi- modal PCN had replaced within-mode edges with edges to other data modes (green). In other words, the multi-modal PCN was able to use features in other modes as “confounding factors” to explain the apparent predictive power of within-mode features. Also, the multi-modal PCN only used ∼6% of all available features (red, **Figure 2E**) whereas the single-mode PCNs (black) often used ∼30% or more. To confirm that improved sparsity and edge replacement was a feature of multi-modality and not simply an artefact of excessive LASSO regularization, we applied the Elimination Algorithm from PGM theory to predict the partial correlation values of single-mode PCNs using the multi-modal PCN (and assuming the data distribution was Gaussian). The partial correlations in single-mode PCNs were predicted at a performance of R=0.92 Pearson correlation (**Figure 2F**), indicating that changes in edge density and removal of spurious edges were theoretically expected. For example, B cell abundance in the Flow PBMC (**Figure 2G**) and blood IFNg levels (**Figure S2**) are instances in which the single-mode PCNs consider features to be substantial direct correlates of both dose and lung CFU, but the multi-modal PCN places these features as intermediates between dose and CFU. Overall, based on both the empirical and the predicted single-mode PCNs to the multi-modal PCNs, we estimated that ∼7-15% of the edges in single-mode PCNs were identified as indirect upon using multi-modal data (**Figure 2H**).

Overall, these observations suggest that the value of multi-modality is not necessarily to provide *more* predictors, but rather to provide the *right* predictors after accounting for confounders, resulting in overall sparser PCNs and eliminating “indirect” correlations by predicting how changes in features propagate through a multi-scale network. This is most notable when comparing the neighbors of dose or Lung CFU between single-mode and multi-modal PCNs (**Figure S2**), where a single-mode PCN for PBMCs identifies multiple cell-types in the blood as direct correlates of dose, but the multi-modal PCN removes most of them (**Figure 2G**), and B cells are similarly dropped from the direct correlates of CFU. In the context of IV-BCG vaccination, the PCN analysis shows that there are many paths traversing multiple modes between dose and CFU, which aligns with the prior knowledge that mechanisms of vaccine-mediated protection involve multiple biological scales.

### Direct, intermediate, and peripheral correlates of vaccine-mediated protection

Having built a sparse network of direct correlations between the immune features, vaccine dose, and Mtb burden measured by CFU, we sought to understand the network paths that propagate the effects of the vaccine to affect CFU, in contrast to paths that stem from vaccine dose but propagate to “off-target” effects, or paths that lead to CFU protection “naturally” independent of vaccination. To determine which of the measured immune features were intermediates along the paths to vaccine-mediated protection, we evaluated the various ways that dose and CFU could be separated from each other on the PCN by removing combinations of nodes, thereby cutting off all paths connecting dose to CFU. Specifically, we determined all the *minimal separator sets* of dose and lung CFU (i.e. any combination of nodes that upon removal would leave no paths connecting dose to CFU, ‘minimal’ in the sense that each node in the set is necessary for disconnecting dose from CFU). We chose to analyze minimal separator sets because the Global Markov Property^23^ establishes that fixing the values of nodes (i.e. conditioning) in a separator set will prevent the propagation of effects between separated nodes (i.e. conditional independence). In this context, we informally define three kinds of *correlates* of vaccine-mediated protection: *direct* correlates that are direct neighbors (i.e. partial correlate) of dose or CFU, *intermediate* correlates that belong to at least one minimal separator set, and *peripheral* correlates that do not belong to any minimal separator set (See **Figure 3A** for a simple example). By virtue of them being connected through some path to dose or CFU, all three kinds of correlates are expected to have non-zero correlation to dose and CFU, but they have distinct roles, e.g. an intermediate correlate that is not direct may still be a key step for establishing vaccine-mediated protection, while a direct but peripheral node to CFU may represent vaccine-independent protection.

**Figure 3.**
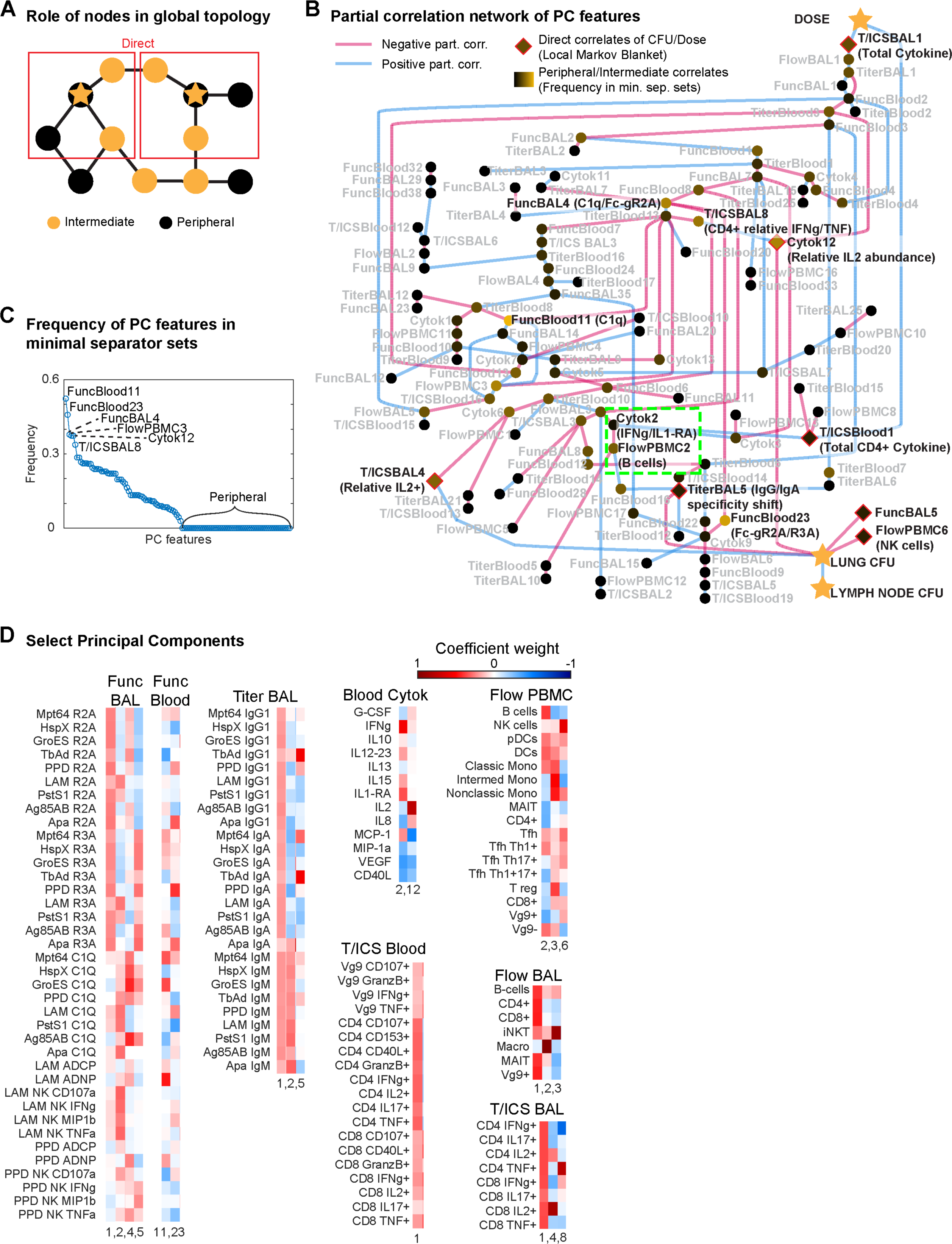
Global graphical analysis of a PCA-PCN. **A)** Schematic example of “direct”, “intermediate” and “peripheral” correlates relative to two starred nodes. **B)** PCA-PCN, with the sign of partial correlations denoted by edge color, direct correlates highlighted in red, and the frequency with which each node appears in all minimal separator sets denoted by node color. Notable regions highlighted in green, and nodes mentioned in text are in bold, black font with a rough interpretation. **C)** Frequency of each node in B in all minimal separator sets. **D)** Weights of original features in select principal components, represented by color.

However, computation of separator sets can quickly become intractable for large and dense networks, so we resorted to making a simpler PCN using latent variables and harsher thresholds. Specifically, we performed Principal Components Analysis (PCA) on each separate data mode, and instead of training a PCN on the original features (e.g. LAM IgM titer in BAL, B cell abundance in PBMC), we trained on the Principal Components (PCs) of each data mode (e.g. Serology Titer in blood PC7, PBMC PC3). We then increased the PCN modeling stability threshold from 80% to 99%, and only kept partial correlations with magnitude >0.15, resulting in the PCA-PCN shown in **Figure 3B** with 115 nodes and 138 edges in the same connected component as dose and CFU. We found that some of the PCs were reasonably interpretable upon examining their weights in the original features (**Figure S3**), e.g. the Serology Titer data modes for both blood and BAL compartments had PC1 showing uniform, positive weights for nearly all antigens and antibody isoforms, which we interpret as total antibody titer, while PC2 had opposite sign weights between the IgMs and IgGs/IgAs, which we interpret as the relative ratio between the two groups of antibodies, possibly corresponding to the degree of antibody class-switching. We then computed the minimal separator sets of the PCA-PCN^25^, finding all 1.26 x 10^6^ minimal separator sets amidst the 2^155^ ∼ 4 x 10^46^ possible subsets, and tabulated the frequency with which each node appeared in these separator sets (**Figure 3C**). Of the 115 PC correlates, 61 were intermediate, and 54 were peripheral. Dose had 2 direct PC correlates and lung CFU had 5 (diamonds in **Figure 3B**).

Reassuringly, the PCA-PCN’s direct correlates overlapped with those originally fitted by the original multi-modal PCN: direct PC-correlates of dose are the T-cell cytokine PC1’s (T/ICSBlood 1 & T/ICSBAL1), which correspond to total CD4 cytokine expression in blood and CD4 or CD8 cytokine expression in BAL (**Figure 3D**), in agreement with the many T-cell cytokines having partial correlation to dose in the original PCN (**Figure 1E)**. The direct PC-correlates of lung CFU include Flow PBMC PC6 (FlowPBMC6), which was effectively NK cell abundance (**Figure 3D**), and BAL titer PC5 (TiterBAL5), which was effectively a shift towards favoring IgGs and IgAs with specific reactivity (**Figure 3D**) to antigens that were all also observed in the multi-modal PCN in **Figure 1E**. Of note, the *relative* abundance of IL2 to other cytokines showed up in the PCA-PCN as CFU-direct, with negative partial correlation for Blood Cytokine PC12 (Cytok12) but positive partial correlation to BAL T-cell cytokine PC4 (T/ICSBAL4; **Figure 3BD**), suggesting that the role of IL2 in protection is compartment-dependent.

Noteworthy examples of intermediate correlates of dose and CFU included PBMC B cell abundance (FlowPBMC2) and IFNg (Cytok2) in the blood (green highlight in **Figure 3B**), as was hinted by the single-mode PCN analysis in **Figure S2**. Notably, Blood Cytokine PC2 included not just IFNg but also IL1-RA. The intermediates found most frequently in minimal separators included the relative amount of CD4+ IFNg to TNF expression (T/ICSBAL8), and general complement component 1q (C1Q) and Fc-gamma receptor IIA and IIIA (R2A, R3A) activity (FuncBlood11 & 23 and FuncBAL PC4) previously as a key predictor of protection^19^. We also note that NK cell abundance (i.e. FlowPBMC6) was a peripheral yet direct correlate of CFU, suggesting that NK cells in the blood contribute “natural” protection that is not vaccine-mediated. More broadly, the PCA-PCN shows that leading PCs (i.e. 1-3) for almost all immune data modes are near the dose node, indicating that vaccination is the major driver of variance observed in each data mode and scale of the immune system. Meanwhile, the intermediate and CFU- direct nodes include many non-leading PCs, suggesting that Mtb-control is governed by subtler variations in the immune system.

### Experimental perturbation of an intermediate node: B cell depletion by Rituximab

We wished to validate our IV-BCG model using data from a separate study, ideally one in which a targeted immune intervention was employed that could lead to effects propagating through the network. Given prior associations between humoral features and IV-BCG protection^14,19^ and that B cells are an intermediate node between dose and CFU in our PCN models, we performed an experimental study depleting B cells prior to and during the initial phase of IV-BCG vaccination, using anti-CD20 antibody (Rituximab) (**Figure 4A**). Our reasoning, based on our previous study, was as follows: a) When delivered via IV, numbers of live BCG are reduced by from 10^7^ to ∼10^3^ within 4 weeks and rarely cultured from vaccinated animals months later, indicating that BCG does not survive very long in IV vaccinated macaques; b) Mycobacterial specific antibodies peak between 4-8 weeks post IV BCG in macaques; and c) If B cells were not present during the early phase of vaccination, antibody production would be lower and when B cells recovered after Rituximab administration ceased, there would be minimal live BCG to provoke a strong antibody response. We also had prior experience with Rituximab in macaques^26^, which indicated that prolonged administration (>10 weeks) of Rituximab was deleterious to some macaques. Thus, we initiated Rituximab in nine rhesus macaques 3 weeks prior to BCG IV and continued every 3 weeks up to 6 weeks post-vaccination (**Figure 4A**). Control animals were administered saline during the vaccination phase at the same intervals as Rituximab. Macaques were challenged with low dose Mtb Erdman via bronchoscope 24 weeks post-BCG IV vaccination, monitored with serial PET CT scans during the challenge period, and necropsied 12 weeks post-Mtb challenge. Four unvaccinated animals were also challenged with Mtb, and we included six historical unvaccinated and Mtb challenged controls as comparators for vaccine success.

**Figure 4.**
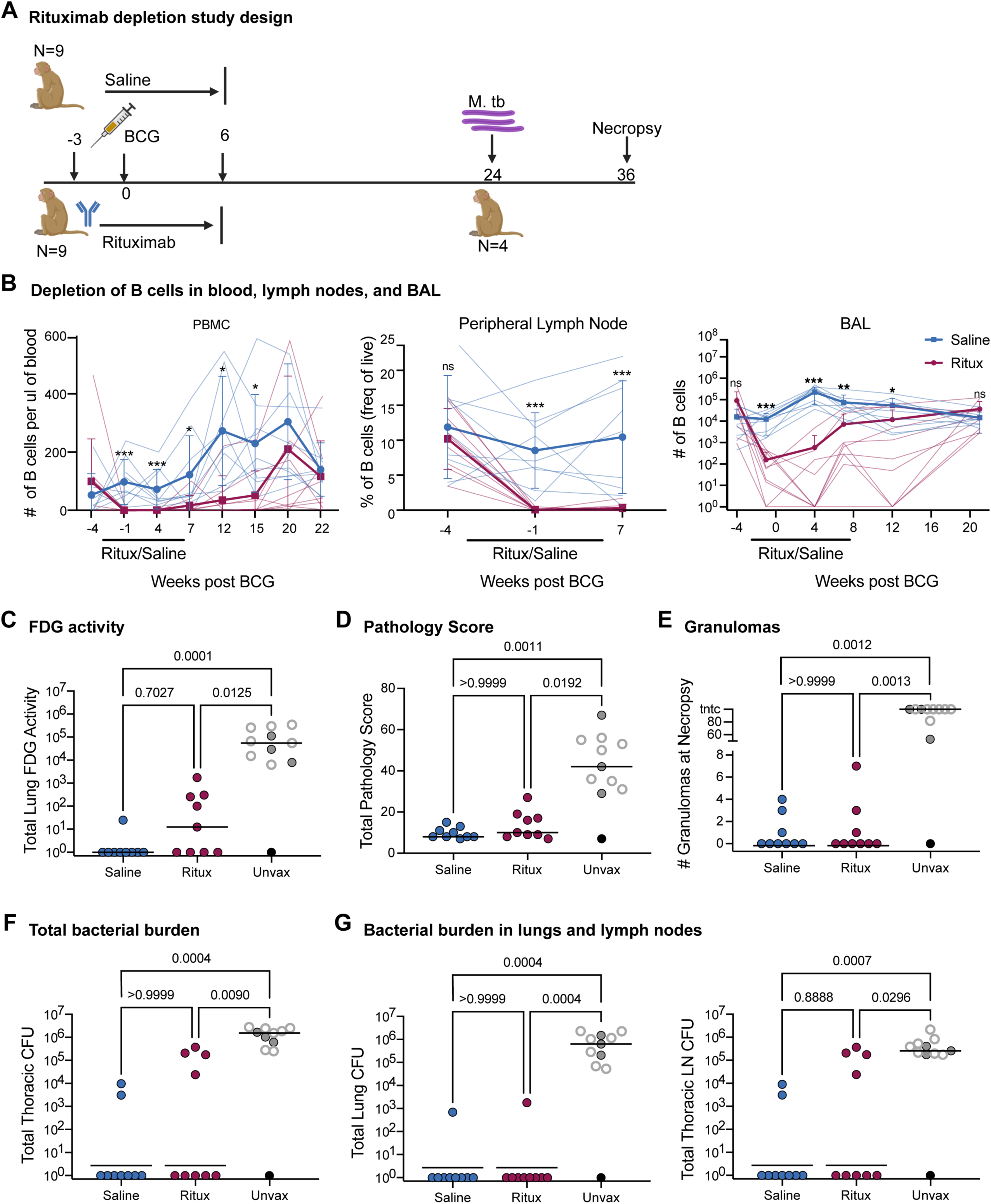
B cell depletion in macaques was successful and did not impair immune responses. **A)** Study design: 9 Rhesus macaques were either treated with saline or Rituximab 4 times, from 3 weeks pre BCG through 6 weeks post BCG. Mtb strain Erdman was administered via bronchoscopes 24 weeks post BCG. Animals were necropsied 12 weeks post-Mtb challenge. Four unvaccinated animals were included as concurrent controls. **B)** B cell depletion due to Ritiximab treatment in blood (# B cells/ul), peripheral lymph nodes (frequency of live cells) and airways (# B cells/BAL). B cells were determined by flow cytometry using CD20 as a B cell marker. Black bar under x axis designates time period when Rituximab was given from 3 weeks before BCG to 6 weeks after BCG vaccination. Rituximab treated animals are shown in blue while saline treated animals are shown in red. Average values for 9 monkeys per group are shown. Mean and SD shown. Multiple Mann-Whitney Tests with Holm-Šídák method adjusted p-values reported: ns - p > 0.05, * - p < 0.05, **, p < 0.01, ***, p < 0.001 **C)** Total PET inflammation as measured by FDG activity in the lung at necropsy. **D)** Total pathology score from necropsy. **E)** Number of granulomas found at the time of necropsy. **F)** Total thoracic CFU at necropsy as counted in tissues. **G)** Respective CFU in lung and thoracic lymph node**. C-G)** Each point is an animal and lines represent medians. Open gray circles are historical controls. Black circle represents clinically uninfected animal. Saline and Ritux groups have 9 animals and the unvaccinated group has 11 animals. Statistics: Kruskal-Wallis test with Dunn’s multiple comparisons test p-values reported.

### Rituximab successfully depleted B cells in blood, BAL and lymph node

Serial blood draws, bronchoalveolar lavage (BAL) and peripheral lymph node biopsies were performed to assess B cell depletion throughout the Rituximab infusions (**Figure 4B**). Baseline levels in blood were measured prior to the start of Rituximab every 2-3 weeks up to 7 weeks, then monthly to assess the level of B cells based on flow cytometry using cell surface markers CD3, CD20, CD79a.

Control group (saline) macaques were treated similarly. B cells were significantly depleted in blood by Rituximab prior to BCG IV administration, and only reached the level of the saline control animals by 20 weeks post-BCG. In lymph nodes, B cells were essentially undetectable in Rituximab-treated macaques at 4 and 7 weeks post-vaccination; lymph node biopsies were not performed past this time point. In the airways, B cells were significantly lower in Rituximab treated macaques up to 12 weeks post-vaccination and were at the level of control animals by 20 weeks post-vaccination. Thus, Rituximab was successful at depleting B cells in blood, tissues, and airways for at least 12 weeks post-vaccination.

### Depletion of B cells did not impair T cell responses in airways

To assess immune responses to BCG IV in the BAL, cells prior to and after BCG vaccination were counted and stained for cell surface markers and intracellular cytokines. Although the number of CD4 and CD8 memory T cells increased substantially after BCG vaccination, and there was an ∼100 fold increase in activated CD4 or CD8 memory T cells making a cytokine (IFNg, TNF, IL2 or IL-17) or a TH1 cytokine (IFNg, TNF or IL2) after 6 hr stimulation with mycobacterial whole cell lysate (WCL), there were no significant differences between the Rituximab and Saline groups (**Figure S4A**). Thus, B cell depletion did not appear to impair induction of T cells expressing cytokines in the BAL.

### Immune responses in lung of vaccinated and B cell depleted animals

To assess whether early B cell depletion affected immune responses in the lungs of vaccinated animals, we excised and processed three random lung lobes from each animal in each group, which were excised and processed into single cell suspensions for flow cytometry. CD3+CD4 and CD3+CD8ab T cell responses were assessed at baseline or after stimulation with Mtb specific peptides ESAT6 and CFP10 or with mycobacterial WCL. As expected, CD4 T cells did not respond to stimulation with ESAT6 and CFP10 but did respond to WCL stimulation in the saline and Rituximab treated groups, though the result did not reach significance in Rituximab-treated animals (**Figure S4B**). In contrast, CD8 T cells did not respond to WCL or ESAT6/CFP10 with cytokine production (**Figure S4C**). There were no statistical differences between total cell counts in lung tissues from vaccinated animals given Rituximab or saline cell surface markers (CD3, CD20, CD79a, CD4, CD8a, CD8b) and no differences were seen in the numbers of T or B lymphocytes in the lung (**Figure S4D**). Thus, following discontinuation of Rituximab 6 weeks after BCG IV vaccination, B cells repopulated the lungs either during vaccination or after Mtb challenge.

### Rituximab did not significantly impair protection against Mtb afforded by BCG IV

Serial PET CT scans using 18F-fluorodeoxyglucose (FDG) were used to assess Mtb infection and disease progression (**Figure 4C**). Total lung FDG activity is a proxy for lung inflammation and is correlated with total thoracic bacterial burden in macaques ^27^. Both BCG IV vaccinated groups had significantly lower total lung FDG activity compared to unvaccinated controls (saline p=0.001 and Rituximab p=0.0125) and there was no significant difference between Saline and Rituximab groups (p=0.7027).

At 12 weeks post-Mtb challenge, detailed necropsies were performed on all animals using the final PET CT scan as a map for obtaining each individual lesion. All granulomas and other Mtb-related pathologies, as well as all thoracic and peripheral lymph nodes, spleen and liver were individually harvested, and single cell suspensions prepared for quantifying bacterial burden and for immunologic analysis. Gross pathology scores indicated that both BCG IV vaccinated groups were significantly lower than the unvaccinated controls, there was no difference between Saline and Rituximab BCG IV vaccinated macaques (**Figure 4D**); and there was also no difference in the number of granulomas found in Rituximab treated vs saline treated BCG IV vaccinated animals (**Figure 4E**). As seen in our previous study, most BCG IV vaccinated animals from both groups had no discernible granulomas^16^. In contrast, unvaccinated animals had significantly higher numbers of granulomas at necropsy compared to either Rituximab or Saline BCG IV vaccinated animals.

Our primary outcome measure was total thoracic CFU, calculated by adding the CFU from each sample from the lung (including granulomas, other pathologies, and grossly uninvolved lungs) and thoracic lymph nodes to obtain the total bacterial burden (**Figure 4F**). Seven of the 9 saline treated vaccinated animals were sterile (i.e. no Mtb recovered) while 5 of the 9 Rituximab treated animals were sterile. Both vaccinated groups had significantly lower total thoracic CFU compared to unvaccinated animals, but there were no significant differences between vaccinated groups in total bacterial burden or when burden was separated into total lung CFU and total lymph node CFU (**Figure 4G**).

### Rituximab treatment significantly reduced mycobacterial specific antibodies in blood and airways

Given the highly efficient depletion of B cells by Rituximab in our system, we next sought to determine whether Rituximab treatment concomitantly depleted the robust plasma and bronchoalveolar lavage fluid (BAL) antibody titers characteristic of IV BCG immunization^17^. This would not be guaranteed as Rituximab does not directly impact CD20-negative antibody-secreting cells, most notably plasma cells which reside in the bone marrow and do not express CD20. Moreover, after cessation of Rituximab treatment and prior to Mtb challenge, it is possible that residual BCG could persist to induce humoral immune responses once CD20+ B cells return. To assess both glycan- and protein-targeting humoral immunity elicited by IV BCG vaccination in the Rituximab- and saline-treated macaques over the course of the study, IgG1, IgA, and IgM antibody titers to lipoarabinomannan (LAM), an abundant Mtb cell wall glycolipid, and culture filtrate proteins from Mtb (CFP) were evaluated by a custom Luminex assay^28^.

As expected, saline-treated macaques exhibited detectable LAM- and CFP-specific antibody titers across antibody isotypes in the plasma (**Figure 5** and **S5**). By contrast, antibody responses in the Rituximab-treated group were markedly weaker, with only 3 macaques generating antigen-specific IgG1 and IgA responses above background, and 2 generating antigen-specific IgM responses above background (**Figure 5A** and **S5A**). Likewise, in the BAL, saline-treated macaques mounted robust IgG1, IgA, and IgM antibody titers following IV BCG vaccination, while glycan- and protein-targeting antibody responses in the Rituximab group were largely muted (**Figure 5B** and **S5B**).

**Figure 5.**
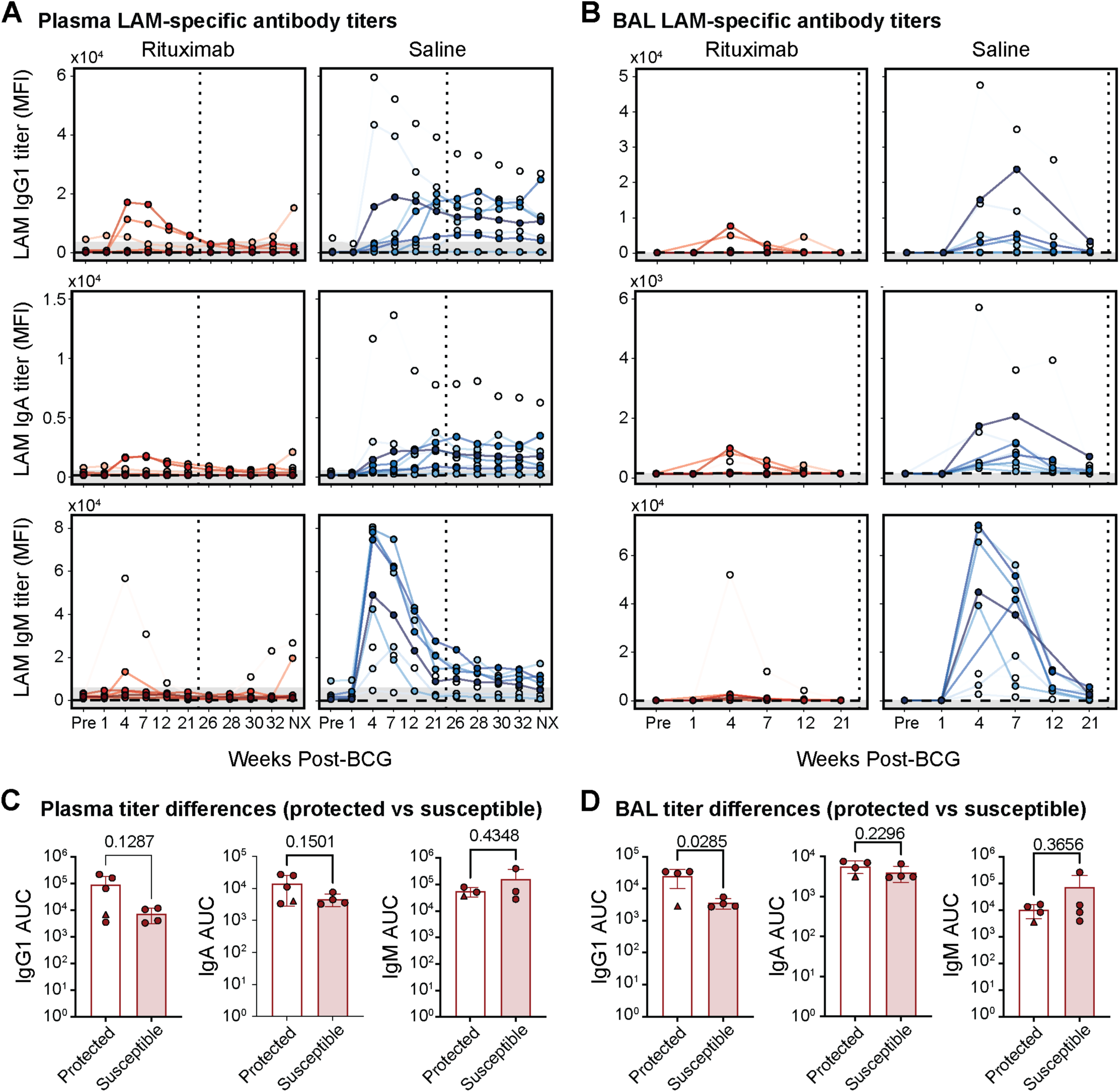
LAM-specific antibody responses. A, **B)** LAM-specific antibody titers in plasma (**A**) and BAL fluid in (**B**). Rituximab- and saline-treated macaques following IV BCG immunization. IgG1 (top), IgA (middle), and IgM (bottom) antibody titers specific to LAM measured via Luminex. Y-axis is median fluorescence intensity (MFI). X-axis shows time in weeks, including pre-vaccination at week -4 (Pre) and necropsy at week 38 (NX). Each point represents the duplicate average from a single macaque at the corresponding timepoint. Rituximab-treated macaques are red and saline-treated macaques are blue. Vertical dotted line indicates the time of challenge (week 23). Horizontal dashed line is the PBS control. Grey shaded area is the background level set to 2 standard deviations above the mean MFI of the pre- vaccination samples. Measurements at least 2 standard deviations above the mean MFI of the pre- vaccination samples are considered above the limit of detection. **C, D**) Comparison of LAM-specific antibody titers between protected (total Mtb CFU = 0) and susceptible (total Mtb CFU > 0) Rituximab- treated macaques in the plasma (**C**) and BAL (**D**). Area under the curve (AUC) values were calculated using longitudinal Luminex MFI data from the following timepoints: Pre-vaccination (Pre), week 4, 7, 12, and 21. Macaque with 0 Mtb CFU, but residual BCG present in a peripheral lymph node (12620) is represented by the triangle marker. Unadjusted p-values resulting from two-tailed Mann-Whitney U tests are indicated.

Given that antibody responses were reduced but not completely abrogated in the Rituximab group, we next examined the relationship between residual antibody levels in Rituximab-treated macaques, and protection following Mtb challenge. No significant differences in plasma IgG1, IgA, or IgM titers were observed between Rituximab-treated animals with sterilizing protection, and those with breakthrough infections (susceptible) (**Figure 5C** and **S5C**). Yet surprisingly, LAM-specific IgG1 titers in the BAL were significantly higher in protected macaques (p=0.0285), indicating that Rituximab-treated macaques able to control Mtb maintained detectable LAM-specific antibody responses in the airways, despite highly efficient B cell depletion (**Figure 5D**). Together, these data demonstrate that B cell depletion by Rituximab treatment from 3 weeks preceding to 6 weeks following IV BCG immunization results in a profound, but not absolute suppression of vaccine-induced humoral immunity.

### Predicting immune system changes from Rituximab perturbation using the multi-modal PCN

Given we measured many complementary immune features from the Rituximab study, such as T-cell cytokine expression and antibody titers in the BAL, we sought to validate our multi-modal PCN by comparing predicted systems-wide effects of perturbing B cell abundance versus empirically measured changes after Rituximab treatment. Specifically, we assumed that the PCN corresponded to a Gaussian Markov Field (GMF)^23^, and then modeled perturbation of nodes as setting the values of those nodes to some fixed quantity (i.e. conditioning, or an ideal intervention^29^), thereby generating predictions for how means and covariances of immune features change. For example, mean CFU corresponds to average Mtb burden, variance in CFU corresponds to the degree of response heterogeneity between macaques, and correlation between CFU and vaccine dose corresponds to vaccine dose-sensitivity of Mtb.

First, we evaluated the Gaussian assumption and experimental reproducibility of the multi-modal PCN’s predictive power using a different dataset of IV-BCG vaccinated macaques from a different “Route” study^16^, in which the IV-BCG vaccine dose was given at a fixed, high dose to 10 monkeys and measured at various initial times post-vaccination for a subset of the immune features that also appeared in the vaccine dose study. In principle, the fixed IV-BCG dose of the Route study corresponded exactly to conditioning on the vaccine dose variable, and so the quality of PCN predictions of the Route study data reflect technical limitations of the data or fitting procedure as opposed to the limitations of modeling perturbations as conditioning. We evaluated whether observed changes between all pairwise correlations (1328 valid pairs) relative to the vaccine dose study were well predicted (we did not evaluate changes in means due to a lack of normalization standards between the two studies), shown in **Figure 6A** with error bars denoting 2 S.E.’s expected from sampling noise. Predictions reached a Pearson correlation of R=0.44 (p<10^-6^), and we estimated that in the best case scenario, a hypothetical perfect model would be expected to achieve R=0.80 due to sampling noise.

**Figure 6.**
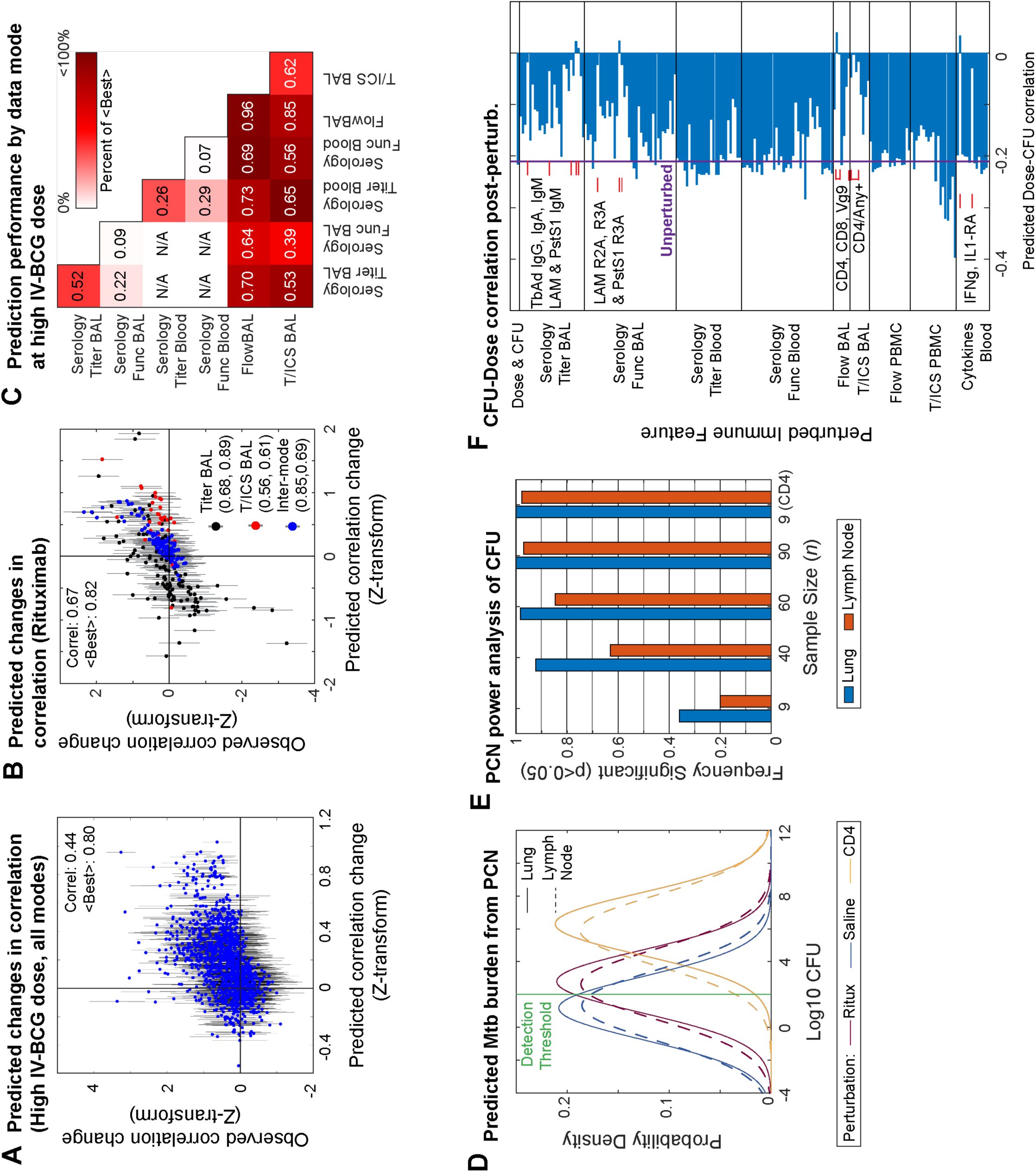
Predicting perturbation outcomes with the multi-modal PCN. **A)** Comparison of predicted versus observed shifts for all pairwise correlations common to both the vaccine dose and Route study, after Fisher’s Z-transformation. Errors bars denote 2 standard errors in either direction, as expected by sampling variation. **B)** Same as A) but for the Rituximab study. **C)** Prediction performance on the Route study based on Pearson correlation (numbers) and relative to the Best expected (color), split by pairs of data modes**. D)** Distributions of CFU in either the lung or lymph node predicted by the multi-modal PCN for the Rituximab and Saline conditions, as well as a hypothetical CD4 depletion. Modeling assumed a practical detection threshold (green) for CFU. **E)** Predicted frequency of detecting a significant difference between Rituximab and Saline conditions for various sample sizes *n*. Predicted frequency also shown for detecting a difference for a hypothetical CD4 depletion at *n=*9. **F)** Predicted correlation between dose and lung CFU upon perturbing any single feature in the multi-modal PCN. Features predicted to bring correlation close to zero are annotated.

Next, we used the multi-modal PCN to model the Rituximab experiment as conditioning on a 100-fold knockdown of the B cell abundance in both BAL and PBMC nodes. We predicted changes in the pairwise correlations (300 valid pairs) relative to Saline with an overall Pearson correlation of R=0.67 (p<10^-6^), relative to an expected best case of R=0.82 for a perfect predictor (**Figure 6B**). Given the comparable performance of the Rituximab predictions to the Route predictions, the ideal intervention assumption seemed to be appropriate for modeling perturbations. Furthermore, we noticed that prediction performance for both datasets varied substantially depending on the data mode (**Figure 6C**), being moderate or poor for Serology measurements compared to a perfect predictor, but quite well when involving flow cytometry data modes, sometimes approaching the expected best level. Overall, the multi- modal PCN was predictively informative despite experimental variation between studies, data limitations, and the Gaussian assumption.

### Statistical power analysis and hypothesis generation using PCN perturbations

In agreement with experiment, the PCN also predicted that the distributions of both lung and LN CFU between the Rituximab and saline conditions would overlap substantially and have only a small difference in means (**Figure 6D**), especially in comparison to a hypothetical perturbation of CD4+ cells in place of B cells. As a form of statistical power analysis, we then used the predicted CFU distributions to roughly estimate the cohort size *n* needed to have a high probability of detecting a statistically significant difference between the saline and Rituximab groups. To account for the fact that experimental CFU distributions are highly non-Gaussian, and that in the original vaccine dose study data we observed that macaques either had 0 lung CFU or >10^2^ Lung CFU but no values in-between (possibly due to detection limits or a step-like non-linearity determining whether Mtb can establish itself or not), we additionally assumed that any samples from the modeled CFU distributions with values <10^2^ would simply show up experimentally as 0. Under this assumption, we computed the probability of significantly detecting a difference between Rituximab and Saline conditions at p<0.05 for various values of *n* in **Figure 6E**. At our cohort size of *n* = 9, the PCN predicted only ∼25% chance of significance and that *n=*40-90 macaques were needed to achieve >90% chance of detecting significant differences in lung or lymph node CFU, which would be unfeasible for macaque studies. In contrast, we found that *n*=9 would be ample to distinguish Saline from the hypothetical CD4 knockdown. Thus, the PCN supports the conclusion that B cells play a subtle role in IV-BCG mediated protection, but would require an infeasible cohort size to confirm with high statistical significance.

Finally, as evinced from our hypothetical CD4 knockdown, we used the PCN to predict how wider perturbations to other immune nodes would affect the correlation between vaccine dose and lung CFU. This could help define features that are important for vaccine-mediated protection. Specifically, whereas the PCN model originally had a dose-CFU correlation of -0.2, we computed the correlation after perturbing any single node (**Figure 6F**), taking note of increases or decreases in correlation. We found that certain perturbations would bring dose-CFU correlation to ∼0, interpreted as loss of crucial intermediates necessary for vaccine-mediated prediction. For example, the PCN found that CD4+, CD8+, and Vg9+ abundance in the BAL were crucial intermediates, alongside expression of any of the measured cytokines in BAL CD4+ cells, and IFNg and IL1-RA cytokines in blood. Various BAL IgM titers, were also crucial, corroborating a recent systems serology study of IV-BCG vaccinated NHPs^17^. These features also appeared in the PCA-PCN analysis as important intermediates. Also corroborating the PCA-PCN analysis, a perturbation at NK cells in blood had limited effect on CFU-dose correlation even though NK cells were a direct neighbor of lung CFU in the multi-modal PCN, consistent with the earlier interpretation of NK cells as a peripheral node providing “natural” protection but not vaccine-mediated protection. This exercise helps prioritize perturbation experiments that could help validate the importance of key mediators of the IV-BCG protective response.

## Discussion

We have applied partial correlation networks (PCN), a type of probabilistic graphical model (PGM), to perform multi-modal data integration of measurements across different scales of the immune system in a cohort of macaques to better understand the immune responses associated with IV-BCG vaccine-mediated protection. In contrast to correlation or covariate analysis previously performed on multimodal data from IV-BCG-vaccinated animals, graphical analysis using PCNs allows us to understand how the influence of individual nodes propagate through the multi-scale network of immune interactions (cellular abundances, antibody titers and function, cytokines), such as how vaccine effects lead to protection against Mtb, off-target effects, or how certain features of protection may be independent of vaccination. Not only were PCNs useful for describing different cascades of effects in the immune system following vaccination, we also showed examples of how PCNs could then be used to infer outcomes of perturbations and perform statistical power analysis.

The combination of PCN modeling with multi-modal data integration was key for learning data-driven models with a topology that begins to broadly resemble the mechanistic topology expected of a complex immune system response to vaccination. First, multi-modal data integration is theoretically necessary given that mechanistic sequences of immune responses often span various scales that cannot all be measured by a single experimental modality. While our models leveraged data modalities from a few scales, the measurements within each scale are not exhaustive and there are also obviously missing scales, e.g. protein or RNA expression data. However, we find that a sparse PCN can broadly recapture the overall immune topologies by removing confounding correlations that often appeared in analysis of single data modes. One limitation of PCNs is that while learned PCN models show broadly reasonable topological features that can help suggest mechanistic hypotheses, the findings of the models are still not necessarily mechanisms in any biophysical sense. Second, PCN modeling is a bare minimum statistical framework to quantitatively model the outcomes of sequences of effects in a network compared to more elaborate and data-hungry modeling approaches like differential equation models that can also account for propagating effects. Although we only implicitly included time-variation in our analysis by treating data from different timepoints as samples from the same probability distribution, other PGM tools could also be used explicitly model time-dependence explicitly, e.g. Markov chains. In general, PCNs and PGMs can be practically accommodating when sample size is limited relative to the complexity and dimensionality of multi-modal data, such as in the case of the IV-BCG studies on macaques and across most experimental systems.

The PCN model was also able to reasonably predict how perturbations of different immune features would propagate to changes across the entire network, in spite of limited training data and a Gaussian assumption. Furthermore, we modeled perturbations as conditional distributions following the concept of ideal interventions^29^, which may only represent asymptotic limits of experimental perturbations. Regardless, it was possible to predict changes in means using GMFs as with other regression models, and using PCNs helped us to further predict variances (i.e. heterogeneity) and correlations (i.e. propagated influences), perform statistical power analysis, and understand global graphical structure. It is possible to replace the PCN approach and GMF assumptions with other functional forms that better match the particular behavior of the IV-BCG macaque system, for example by discrete distributions (e.g. Ising model) as opposed to Gaussians. Fitting directed graphical models like Bayesian Networks^30^ as opposed to Markov Fields could also have benefits: while both types of PGMs represent conditional dependencies inherent to a distribution (i.e. of data points), specific types of conditional dependencies can sometimes only be represented graphically by either a DAG or a Markov Field, which might reveal additional insight about interactions. However, under a Gaussian assumption, it is practically guaranteed that a Markov Field already represents all possible conditional dependencies^31^, and we found that GMFs provide substantial computational and theoretical convenience, and that predictions were already of sufficient quality to possibly guide experimental design.

We note that many biological features were interpreted from the PCN by performing global graphical analysis to implicitly understand intermediate nodes, e.g. marginalizing by data modalities, computing minimal separator sets, or examining conditional distributions. In previous applications of data-driven PGMs on biological big data, it was more common to interpret direct neighbors of certain nodes without advancing to global analysis; this may be due to computational infeasibility or difficulty finding rigorous and explicit interpretations for the global features of PGMs, e.g. individual paths in a PGM do not in general have a simple meaning. For example, our choice of analyzing separator sets was motivated by the definition of the Global Markov Property of Markov Fields; analyzing collections of cliques may also be useful, motivated by the factorizations defined by the Hammersley-Clifford Theorem^10^. Intermediates are obviously vital to understanding cascades in a network, and so applications of PGMs to biological big data would benefit from future developments on interpreting and computing global features of PGMs.

The in vivo studies in the BCG-IV vaccinated macaques were intended to determine whether antibodies were necessary for protection against tuberculosis. To accomplish this, B cells were depleted during the peak of BCG IV vaccination when antibodies are being generated. After Rituximab administration was stopped, B cells slowly repopulated in the macaques but anti-mycobacterial antibody levels stayed low compared to control animals. However, the macaque study did not point to a clear requirement for antibodies in BCG IV induced protection against TB. Our data in NHP did suggest that in the B cell depleted group, those animals with somewhat higher antibody levels against LAM appeared to be better protected, suggesting that antibodies may play a minor but still important role in BCG IV-induced protection. However, the PCN predicts that we would need to use many more animals in this type of study to achieve statistically significant differences in infection outcome between vaccinated and Rituximab treated or control animals. Moreover, the PCNs we have constructed suggest that B cells and antibodies do lie along the network paths connecting IV-BCG vaccination to eventual Mtb burden, but that there are many other paths representing other mechanisms that can compensate for the depletion of B cells. While an exhaustive description of all the compensatory paths suggested by our PCN model is not straightforward or easily interpretable, one might speculate that B cells themselves may be playing a role beyond production of antibodies, such as cytokine production, since the B cells do repopulate in the blood and tissues following cessation of Rituximab infusions. Thus, B cells and antibodies may have a minor but still important role in IV BCG induced protection against TB, although it is likely that the strong T cell responses induced by BCG IV^32^ overwhelm most contributions from antibodies. Another possibility is that natural exposure of NHP to environmental mycobacteria prior to BCG vaccination (due to housing conditions, including outdoor rearing) leads to plasma cells making antibodies against these microbes that can cross-react with BCG or Mtb, it is possible that these pre-existing antibodies are sufficient to contribute to BCG-IV induced protection. This is possible because Rituximab does not deplete plasma cells, so such cells and the antibodies produced by them would still exist in Rituximab treated animals. It should be noted that other TB vaccine strategies may have stronger or weaker requirements for B cells and antibodies for protection.

Broadly, we found PCNs to be useful for describing and predicting the propagating effects through multi-scale immune networks, particularly in settings with limited data. Other forms of PGMs, such as Bayesian DAGs, or other distributional assumptions beyond GMFs, might also improve performance. As multi-modal experiments develop to measure more aspects and scales of the intricate networks of biological systems, PCNs and PGMs in general offer a practical way to integrate data to understand multi-scale phenomena in a data-driven fashion.

## Supporting information

Suuplemental Table 1

## Resource availability

### Lead contact

Further information and requests for resources and reagents should be directed to and will be fulfilled by Douglas A Lauffenburger.

### Materials availability

This study did not generate new unique reagents.

### Data and code availability

- This paper analyzes existing, publicly available data. In addition, the Rituximab depletion study data have been deposited at https://fairdomhub.org/studies/1238 and are publically available as of the date of publication.
- All original code has been deposited at Zenodo (DOI:10.5281/zenodo.12604554) and is publicly available as of the date of publication. DOIs are listed in the key resources table.
- Any additional information required to reanalyze the data reported in this paper is available from the lead contact upon request.

## Acknowledgements

This project has been funded in in part with federal funds from the National Institute of Allergy & Infectious Diseases, National Institutes of Health, Department of Health & Human Services, under contract #75N93019C00071 (SMF, JLF, DAL), grant AI150171 (EBI), and grant AI167899 (DAL). We gratefully acknowledge the dedication and contributions of all the veterinary and research technicians at the University of Pittsburgh School of Medicine, along with the contributions of Alex White and L James Frye for help with PET CT scan acquisition and analyses.

## Author Contributions

- Conceptualization – SW, MCC, SMF, JLF, DAL
- Methodology – SW, AJM, EBI, PLL, GA, SMF, DAL
- Software – SW
- Validation – SW, PM
- Formal Analysis – SW, AJM, EBI, PM, HJB
- Investigation –SW, AJM, EBI, PM, MAR, JT, KK, HJB, PLL, JLF
- Resources – CW, PLL, PAD, RAS, MR
- Data Curation – AJM, CW, PM, PLL
- Writing – Original Draft – SW, AJM, JLF, DAL
- Writing – Review & Editing – SW, AJM, EBI, CW, MCC, DM, PLL, JLF, DAL
- Visualization – SW, AJM, EBI, PM
- Supervision – GA, SMF, JLF, DAL
- Project Administration – MCC, DM, CAS, SMF, JLF, DAL
- Funding Acquisition – EBI, SMF, JLF, DAL

## STAR Methods

### Experimental model and study participant details

#### Animals

Adult rhesus macaques (*Macaca mulatta*) (N=22) (10 Female and 12 Male, 3-6 kg, 2-7 yr old) were obtained from BioQual and PrimGen. Details and data for each animal are in **Supplementary Table 1**. Animals were co-housed under BSL2+ conditions during the vaccination phase and in the BSL3 facility during the Mtb challenge phase. Animals were monitored twice daily by our veterinary technical staff and husbandry staff. Enhanced enrichment was provided throughout the study. All procedures were approved by the University of Pittsburgh’s Institutional Animal Care and Use Committee (IACUC).

### Method details

#### Rituximab or Saline Infusion

Macaques were sedated with ketamine and given acetaminophen 80mg and diphenhydramine 1mg/kg based on animal weight prior to infusion. Animals were randomized into 2 groups within 2 cohorts and given either saline or Rituximab (Genentech; purchased from the University of Pittsburgh School of Medicine pharmacy). Four unvaccinated and untreated macaques were included as controls. Dose (50mg/kg) was calculated based on the Dosing Calculator at website: (use 1000 mg/m^2^) as BSA Based Dose; https://reference.medscape.com/drug/rituxan-truxima-rituximab-342243). Rituximab or saline were administered via infusion at 3 weeks prior to BCG inoculation, 2 days prior to BCG, then at 3 and 6 weeks post-BCG. Infusions were performed over 20-30 minutes for each animal and animals were monitored during and post-infusion for any adverse reactions.

#### BCG vaccination

BCG Danish strain 1331 (Statens Serum Institute, Copenhagen, Denmark) was expanded, frozen and received from NIH/Aeras. A vial of BCG (SSI strain) with 3.04 x 10^8^ CFU/ml was thawed and diluted in PBS/Tween 80, briefly vortexed and administered intravenously in 2ml for a dose of ∼5 x 10^7^ per animal.

#### Mtb challenge

Rhesus macaques were challenged by bronchoscope with 10-19 CFU barcoded Mtb strain Erdman as previously described ^34^ 6 months post BCG vaccination; unvaccinated control animals were challenged with a similar dose. The animals were clinically monitored throughout the course of the experiment including physical exams, CBC, erythrocyte sedimentation rate, appetite, behavior and activity. PET/CT was performed at 4 and 8 weeks post Mtb and prior to necropsy (12 weeks p.i.).

#### PET-CT scan and analysis

PET-CT imaging was performed using a Mediso MultiScan LFER 150 integrated preclinical PET CT ^35^. Animals were injected with a glucose analog PET tracer, 2-deoxy-2-(^18^F)Fluoro-D-glucose (FDG), to track overall inflammation ^36,37^. After tracer injection, a period of 50 minutes was observed prior to obtaining PET scans to allow the FDG to be taken up by active tissue. During this time, animals were intubated, anesthetized, and attached to a mechanical ventilator. CT scans were also obtained during the uptake period. During CT acquisition (about 40 seconds in duration), the animal’s breath was held via the ventilator to ensure a clear image of the lungs. Scans were obtained following Mtb infection at 4, 8, and 12 weeks.

All imaging was performed according to biosafety and radiation safety requirements within the Biosafety Level 3 (BSL3) facility at the University of Pittsburgh. Scans were analyzed using OsiriX DICOM viewer^38^ by in-house trained PET-CT analysts ^39–41^.

#### Blood, BAL and tissue processing

Plasma was removed by centrifugation of whole blood at 2500rpm for 15 minutes. Aliquots were frozen for later study. Remaining blood was diluted with 1xPBS and overlaid with Ficoll-Paque PLUS (GE Healthcare) to further isolate PBMC. PBMC were used for flow cytometry while plasma was frozen and shipped to MIT (Alter lab) for systems serology. Bronchoalveolar lavage was performed by instilling and retrieving 4 x 10 ml saline via bronchoscope into a lower lung lobe. BAL fluid was centrifuged, washed with 1x PBS and counted with trypan blue exclusion. BAL cells were analyzed by flow cytometry and fluid was frozen and shipped to MIT for systems serology. Peripheral (axillary) lymph node biopsies were performed pre infusion and 2 and 10 weeks post infusion of Rituximab or Saline to measure depletion of B cells by flow cytometry with CD20 and CD79a cell surface antibodies.

#### Necropsy

Macaques were humanely euthanized at the planned experimental endpoint 12 weeks post Mtb infection. A comprehensive necropsy was performed on each animal, with gross pathology scoring as previously described ^27^. Using the final PET CT scan as a map for lesions, each individual lung granuloma, other lung pathologies, all thoracic lymph nodes, and random sampling (50%) of each lung lobe, spleen and liver were obtained for flow cytometry and bacterial burden determination. Single cell suspensions (either by GentleMacs™ for lung lobes and granulomas or manually through a cell strainer (0.7 um for lymph nodes, spleen and liver) were prepared from each sample and plated individually on 7H11 agar plates for CFU. Plates were incubated for 3 weeks in 5% CO2 incubator and then counted. Actual bacterial burdens per sample were calculated and total thoracic, total lung and total LN bacterial burden calculated for each animal as described previously ^27^.

#### Flow cytometry

After single cell suspensions (necropsy tissue samples, lymph node biopsies, BAL or PBMC) were counted, 0.2-1 x10^6^ cells were resuspended in RPMI/10% Human Serum/L-glutamine/HEPES in a U bottom cell plate. For BAL and Lung and Lymph node cells from necropsy, 50ul of appropriate stimulator (Whole Cell Lysate (WCL) 1mg/ml, ESAT6/CFP10 1mg/ml, Phorbol 12,13-Dibutyrate (PDBU, Sigma) + Ionomycin (Sigma) or media alone were added for 2 hours at 37C. Brefeldin A (GolgiPlug, BD) was added for an additional 4 hours at 37C for BAL and 16 hours for necropsy samples. PBMC and lymph node cells from biopsy were not stimulated or stained for cytokines. Cells were then transferred to a V bottom plate, spun down at 2000 RPM for 3 minutes, washed in 1 x PBS 2x and resuspended in viability dye (Invitrogen) and incubated for 15 minutes at room temperature (RT). Cells were washed in FACS buffer (1xPBS+1% FBS) then stained with the following cell surface antibodies for 30 minutes at 4C (**SI Tables 3, 4, 5)**. PBMC and lymph node cells (from biopsies) were washed again in BACS buffer and fixed with 1% Paraformaldehyde (PFA, Sigma). Tissue from necropsy or BAL were fixed with 100ul Fix/Perm (BD) for 10 min at RT. Cells were washed 2x with Permwash buffer (BD) and stained intracellularly for 30 min at 4C with the following markers (**SI Tables 3, 4, 5**). Data acquisition was performed using an Aurora (Cytek) and analyzed using FlowJo software v10.8.1 (BD, **Figures S6 and S7**).

#### Systems serology

Antibody levels were measured (**Supplementary Table 2)** as previously described^17^. Briefly, LAM and culture filtrate protein (CFP) were coupled to magnetic Luminex beads by DMTMM modification^42^, and carbodiimide-NHS ester coupling^28^, respectively. Coupled beads were incubated with 5uL of plasma or BAL overnight at 4C in 384-well plates (Greiner Bio-One) using the following dilutions: plasma IgG1 (1:30), plasma IgA (1:30), and plasma IgM (1:150). BAL was not diluted. After overnight incubation, the plates were washed and mouse anti-rhesus IgG1 (clone 7H11), IgA (clone 9B9), or IgM (Life Diagnostics, clone 2C11-1-5) antibody was added and incubated shaking at room temperature for 1 hour. Anti-rhesus IgG1 and IgA were from the National Institutes of Health Nonhuman Primate Reagent Resource. The plates were then washed and phycoerythrin (PE)-conjugated goat anti-mouse IgG was added (ThermoFisher, 31861) and incubated shaking at room temperature for 1 hour. Finally, plates were washed and relative antibody levels (PE MFI values) were measured using a FlexMap 3D (Luminex).

Samples were measured in duplicate.

### Quantification and statistical analysis

#### Variable selection from Dose study

We first selected the following groups of features from the variable dose study dataset: IgG1, IgA, and IgM titers for any antigen, all measured antibody functional assays, all flow cytometry measurements of immune cell-type abundances without reference to cytokine staining, and those measuring the proportion of immune cells expressing any individual cytokine (i.e. marginal expression), and all the bulk cytokine measurements in the Blood. We then additionally removed features for which too many data entries were missing, there was limited signal beyond background measurement noise, or when features demonstrated near-exact collinearity with other features. Specifically, in the data modes below, we removed the following features.

Flow BAL: Granulocytes, mDC, NK, pDC

T/ICS BAL: CD4+/IL21+, CD8+/IL21+

FlowPBMC: mDC, Monocytes, T-cells, mMAITs, mCD4+, mCD8+, gdT-cells, mVg9+, mVg9-Blood Cytokines: GM-CSF, IL-17A, IL-18, IL-1b, IL-4, IL-5, IL-6, MIP-1b, TGFa, TNFa

Additionally, we treated different timepoints for any given macaque as different samples from the distribution of the immune network, using all the timepoints for which the aforementioned variables were mostly completely measured for each sample, leaving us with timepoints 0, 4, 8 weeks. Note that in the Systems Serology measurements, there was a timepoint labeled as 9 weeks instead of 8, which we treated as the same timepoint. Also, for Blood Cytokine data, measurements stopped after 4 weeks, but since measurement values over earlier timepoints (6 hours, 2 days, 1 week, 2 weeks) appeared to reach a steady-state before the 4 week timepoint, we made the assumption that Blood Cytokine values at 8 weeks were the same as at 4 weeks. In support of this steady-state timescale being more general, we note that another recent study showed measured cytokine levels reaching steady-state a few days post-vaccination, for various antiviral vaccines in clinical trials^33^.

#### Graphical LASSO and stability selection

For the multi-modal PCN, data were first normalized by applying the arcsinh transformation to all measured features, and then z-scoring each feature whilst ignoring missing entries. Then, we applied a MATLAB implementation of Graphical LASSO^20^ to multiple random subsets of the normalized data to perform stability selection^22^. Specifically, we chose 8 evenly spaced values of the penalty parameter lambda in [0.2 , 1], and for each lambda-value we applied Graphical LASSO to 100 different random 80%-subsets of the data, inputting the sample covariance matrix computed using only samples with no missing entries. We recorded the frequency with which each pairwise edge between features had non-zero magnitude (>10^-6^), and further recorded the maximum frequency of each edge over the range of lambda- values. We then constructed the Partial Correlation Network (PCN) by first choosing the edges with maximum frequency *f* > 0.80, fitting the maximum-likelihood estimate (MLE) Gaussian to the full dataset given those edges^23^, constructing a partial correlation matrix by inverting and rescaling the MLE inverse covariance matrix, and setting all partial correlation values *j* with magnitude *j* <10^-3^ to 0 to define a PCN.

For single-mode PCNs, we performed the same procedure as for the multi-modal PCN, restricted to the features belonging to a single data mode and lung CFU, LN CFU, and dose.

For the PCA-PCN, we performed the same procedure as for the multi-modal PCN, except for two steps. First, we performed Principal Components Analysis (PCA) on each data mode separately (excluding dose, lung CFU, and LN CFU) after the arcsinh and z-scoring, and further z-scored the PC-coordinates of data points before applying Graphical LASSO. Second, we changed the maximum frequency cutoff to *f* > 0.98 and the partial correlation cutoff to *j* > 0.15.

Code for performing stability selection on Graphical LASSO and MLE for a given network topology can be found at https://github.com/shuwang543/ivbcg_partial_correlation_network.

#### Predicting single-mode PCNs using the multi-modal PCN

To predict the single-mode PCNs using the multi-modal PCN, we followed the standard procedure for computing marginal distributions of a Gaussian Markov Field. Specifically, we took the submatrix of the multi-modal PCN’s correlation matrix corresponding to the nodes of a given single-mode PCN, and inverted the submatrix to produce a predicted single-mode precision matrix. We then rescaled the single-mode precision matrix to produce the single-mode partial correlation matrix.

#### Minimal separator sets of a Markov Random Field

We implemented an algorithm for finding minimal separator sets in a graph, as described in by Shen and Liang^25^, using MATLAB. Our implementation can be found at https://github.com/shuwang543/ivbcg_partial_correlation_network.

#### Statistical tests used in Rituximab study

Non-human primate outcome data was tested for normality using the Shapiro-Wilk test. For serial outcome data, multiple Mann-Whitney tests were run comparing saline and Rituximab groups at each time point and Holm-Šídák method adjusted p-values are reported. Necropsy outcome data were analyzed using a Kruskal-Wallis test with Dunn’s multiple comparisons adjusted p-values reported. When comparing stimulation responses of CD4 and CD8 memory cells in lung tissue, Friedman test was used and Dunn’s multiple comparison adjusted p-values are reported. Total cell counts in lungs were compared only in the Saline and Rituximab groups using unpaired t-tests. NHP outcome data was graphed and analyzed in GraphPad Prism 10 for macOS (version 10.1.1).

#### Predicting/inferring perturbation outcomes

To make predictions about perturbations to any subset of nodes, we computed the conditional distribution for the multi-modal PCN after fixing each node *xi* in the subset to a fixed value *ai*, according to the standard procedure for a Gaussian Markov Field^23^. For inferences of single-node perturbations in **Figure 6**, we set *a =* -1. For inferences of the Route study in **Figure 6**, we set Dose = 1. For inferences in the Rituximab study, we set (Dose, BAL #B cells, PBMC #B cells) = (1,-6.3,-1.2) for the Rituximab condition, (Dose, BAL #CD4, PBMC #CD4) = (1,-6.3,-1.2) for the hypothetical CD4 perturbation, and Dose = 2 for the Saline control. Perturbation values were chosen to equal the empirical changes at each node from the corresponding experiments, accounting for the arcsinh transformation and z-score rescaling performed on the data as preprocessing.

#### Estimating the Best expected performance of a perfect predictor

We defined the Best expected performance of a perfect predictor to be the square-root of the proportion of total observed variance *σ*^2^*_tot_* leftover after subtracting sampling noise variance *σ*^2^*_samp_*. We estimated *σ*^2^*_samp_* as the average value of 1/(*N_i_*− 3) over *i* for all the different pairwise correlations, for *N_i_* the number of samples for which neither of the two features in a pair were missing data entries, i.e.

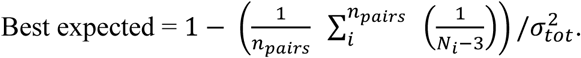

#### Statistical power analysis of CFU differences between conditions

To roughly estimate the probability of observing significant differences in lung or lymph node CFU for a given sample size *n* between Saline and Rituximab/CD4 perturbation conditions, we defined the sampling distributions of either lung or lymph node CFU for each condition as the corresponding conditional distribution of the multi-modal PCN used above for prediction/inference, marginalized to either lung or lymph node CFU. Since each sampling distribution defined in this manner was Gaussian, but observed CFU distributions from the Dose study were highly non-Gaussian, samples from these CFU distributions were defined as 0 if their CFU-value was below a threshold; we chose 10^2^ as the threshold to imitate the apparent threshold observed in the variable dose study. Using this sampling scheme, we took *n* samples from each CFU distribution, and performed the Mann-Whitney U-test to test for significant differences at a significance level of 0.05; we repeated this 300 times for any given *n*, and recorded the frequency of significant tests.

### Key resources table

| REAGENT or RESOURCE | SOURCE | IDENTIFIER |
| --- | --- | --- |
| Antibodies |  |  |
| Antibodies in Rituximab depletion study | See: https://fairdomhub.org/studies/1238 |  |
| Bacterial and virus strains |  |  |
| IV-BCG vaccine | Statens Serum Institute, Copenhagen, Denmark | BCG Danish Strain 1331 |
| Mycobacterium tuberculosis strain | See: https://fairdomhub.org/studies/1238 |  |
| Chemicals, peptides, and recombinant proteins |  |  |
| Rituximab | Genentech, Inc. | NDC 50242-053-06 |
| Deposited data |  |  |
| Data from BCG route study | https://doi.org/10.1101/2023.07.31.551245 | https://fairdomhub.org/studies/1198 |
| Data from IV-BCG dose study | https://doi.org/10.1016/j.chom.2023.05.006 | https://doi.org/10.1016/j.chom.2023.05.006 |
| Flow cytometry cell counts and systems serology data for Rituximab depletion study | This paper | https://fairdomhub.org/studies/1238 |
| Experimental models: Organisms/strains |  |  |
| NHP Macaques | See: https://fairdomhub.org/studies/1238 |  |

| Software and algorithms |  |  |
| --- | --- | --- |
| MATLAB R2022a | Mathworks Inc. | https://www.mathworks.com/ |
| MATLAB implementation of minimal separator set enumeration algorithm | This paper; Shen and Liang<sup>25</sup> | https://github.com/shuwang543/ivbcg_partial_correlation_network<br>DOI:10.5281/zenodo.12604554 |
| MATLAB implementation of Graphical LASSO | Chen<sup>20</sup>; Friedman et al. <sup>21</sup> | https://github.com/xiaohuichen88/Graphical-Lasso |

**Supplemental Table 1. Rituximab B-cell depletion study animal details.**

**Supplemental Table 2. Rituximab B-cell depletion study systems serology data.**

---

**Figure S1.**
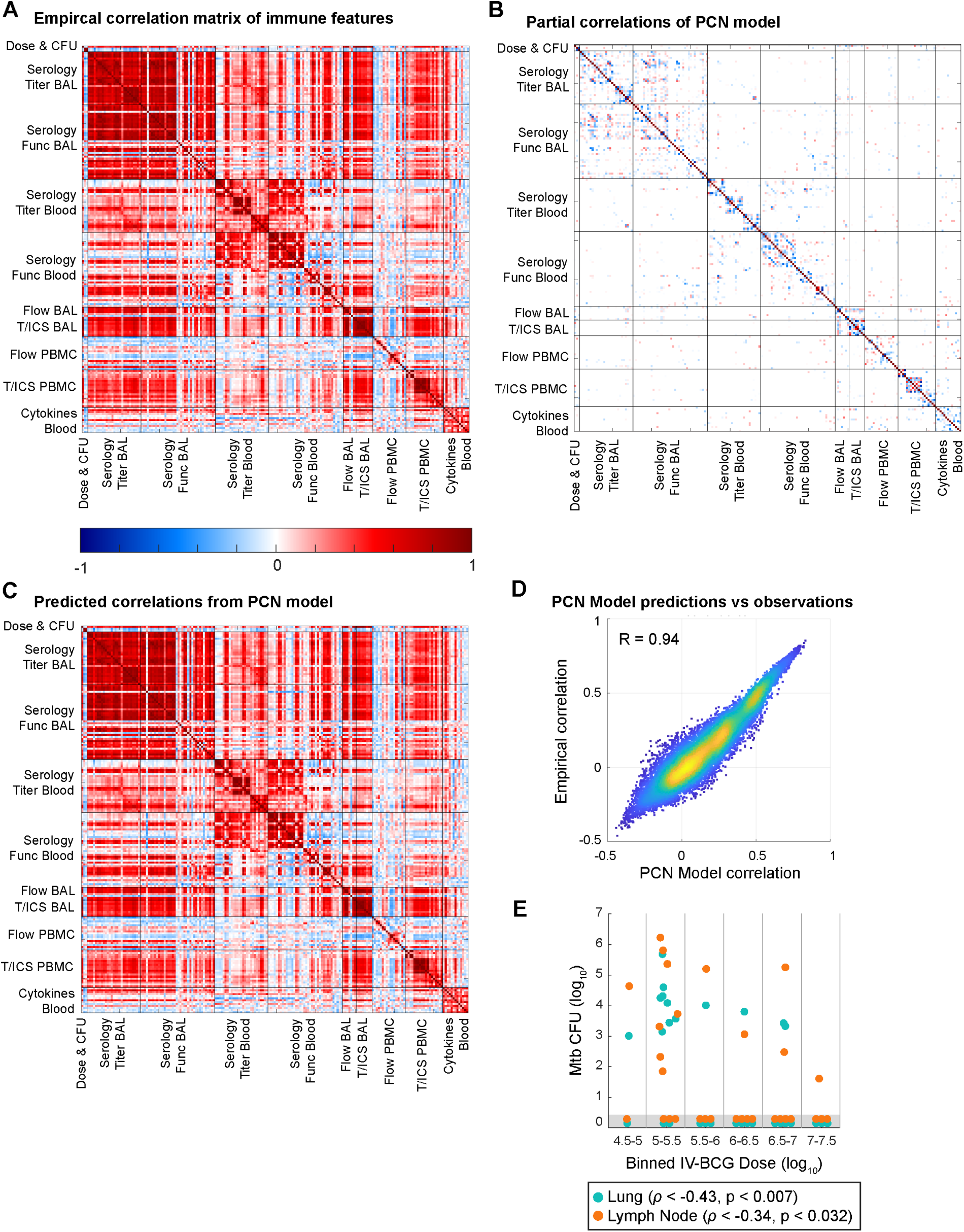
Partial correlation network structure and predictions for IV-BCG vaccinated macaques. **A)** Observed correlation matrix between all immune features. **B)** Partial correlation matrix of the PCN. **C)** Predicted correlation matrix from PCN. **D)** Comparison of correlations observed empirically from the data versus predicted by the PCN. Entries are only included if there was no edge in the PCN corresponding to that pair. **E)** Dependence of lung and lymph node CFU on IV-BCG dose. Spearman correlation, one-tailed.

**Figure S2.**
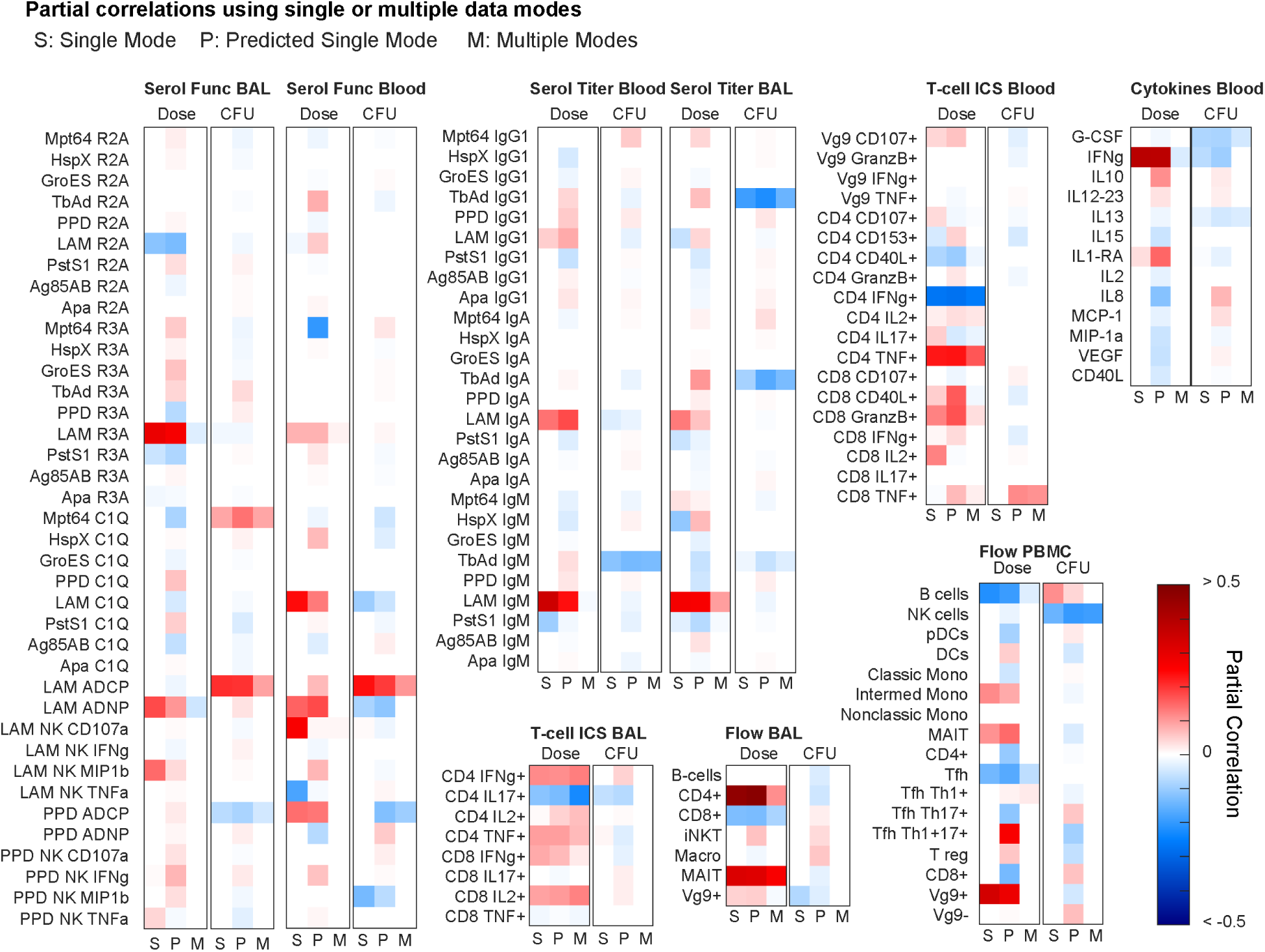
Comparison of partial correlations for single- and multi-modal PCNs. Same as Figure 2G, but for each data mode.

**Figure S3.**
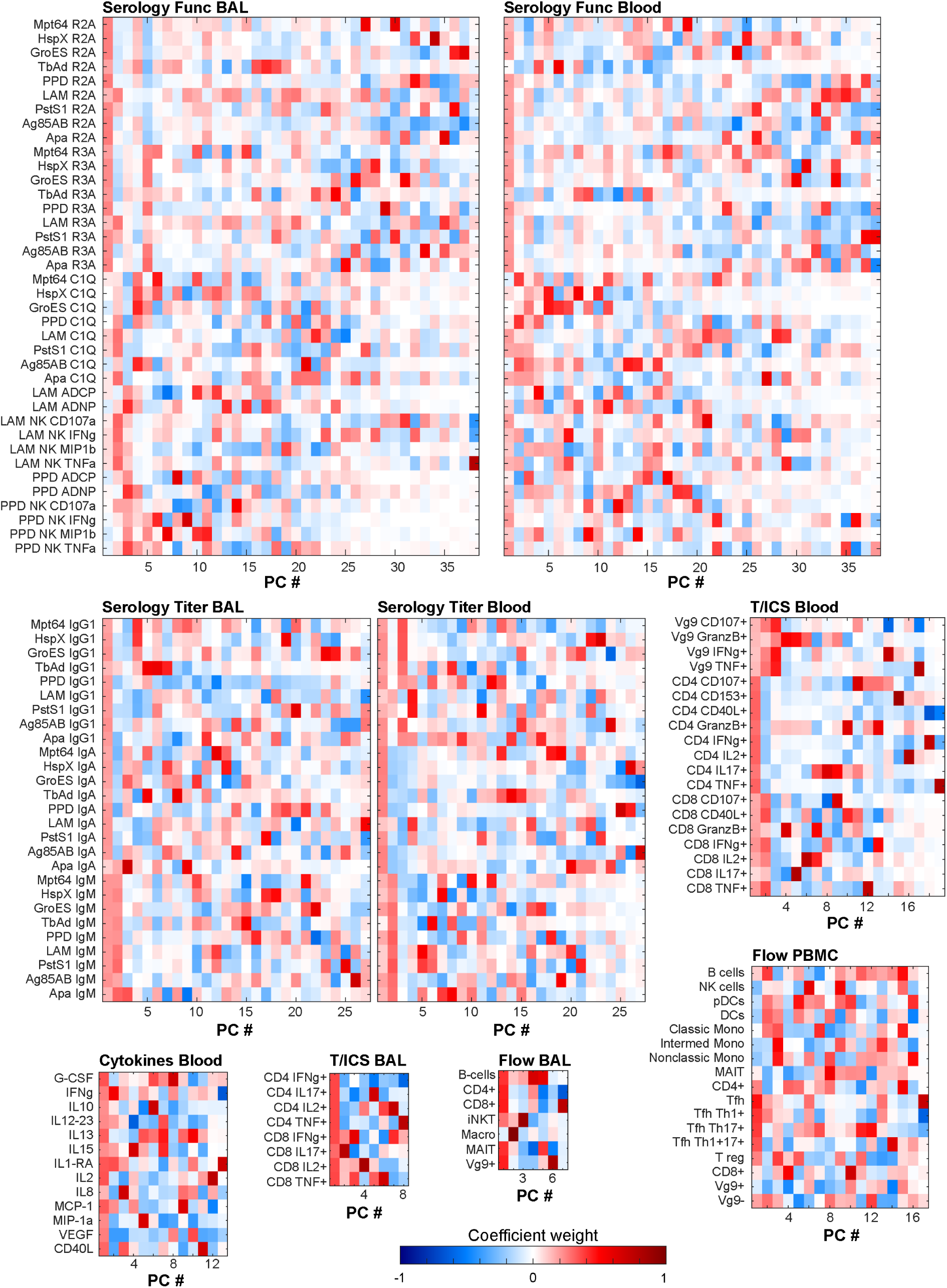
Principal components of single data modes. Weights of original features of principal components for each data mode.

**Figure S4.**
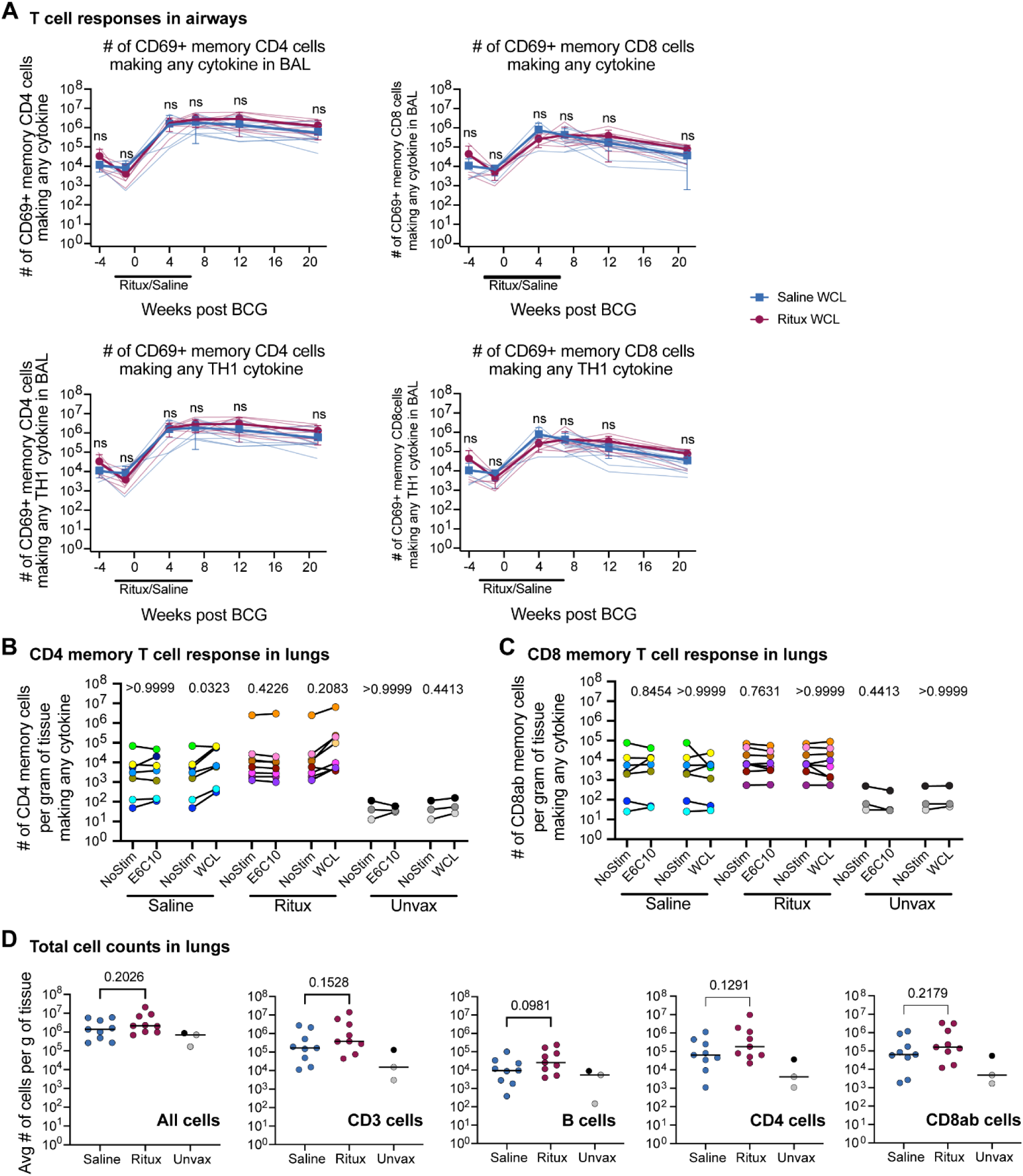
T cell responses upon B cell depletion. **A)** Number of CD4 memory cells in BAL using CD45RA and CD28 to differentiate memory cells via flow cytometry. Rituximab treated animals are shown in blue while saline treated animals are shown in red. Mean and SD shown. Statistics: Multiple Mann-Whitney Tests with Holm-Šídák method adjusted p-values reported: ns - p > 0.05. **B, C)** CD4 and CD8ab memory T cells making any cytokine (IFNgamma, IL-2, TNFalpha or IL-17) in the lung at time of necropsy. Cells were unstimulated or stimulated with ESAT-6/CFP10 peptides or Whole Cell Lysate (WCL). Each point (colored by animal) represents a median from 3 separate lung lobes from each animal. Saline group consists of 7 animals, Ritux group 8, and Unvax group 3. Statistics: Friedman test (for paired data) with Dunn’s multiple comparison adjusted p-values reported. **D)** Cell counts from hemacytometer count of single cell suspensions at time of necropsy and as measured from cell surface markers during flow cytometry. Statistics: Unpaired t test p-values shown. **B-D)** Black circle represents clinically uninfected animal.

**Figure S5.**
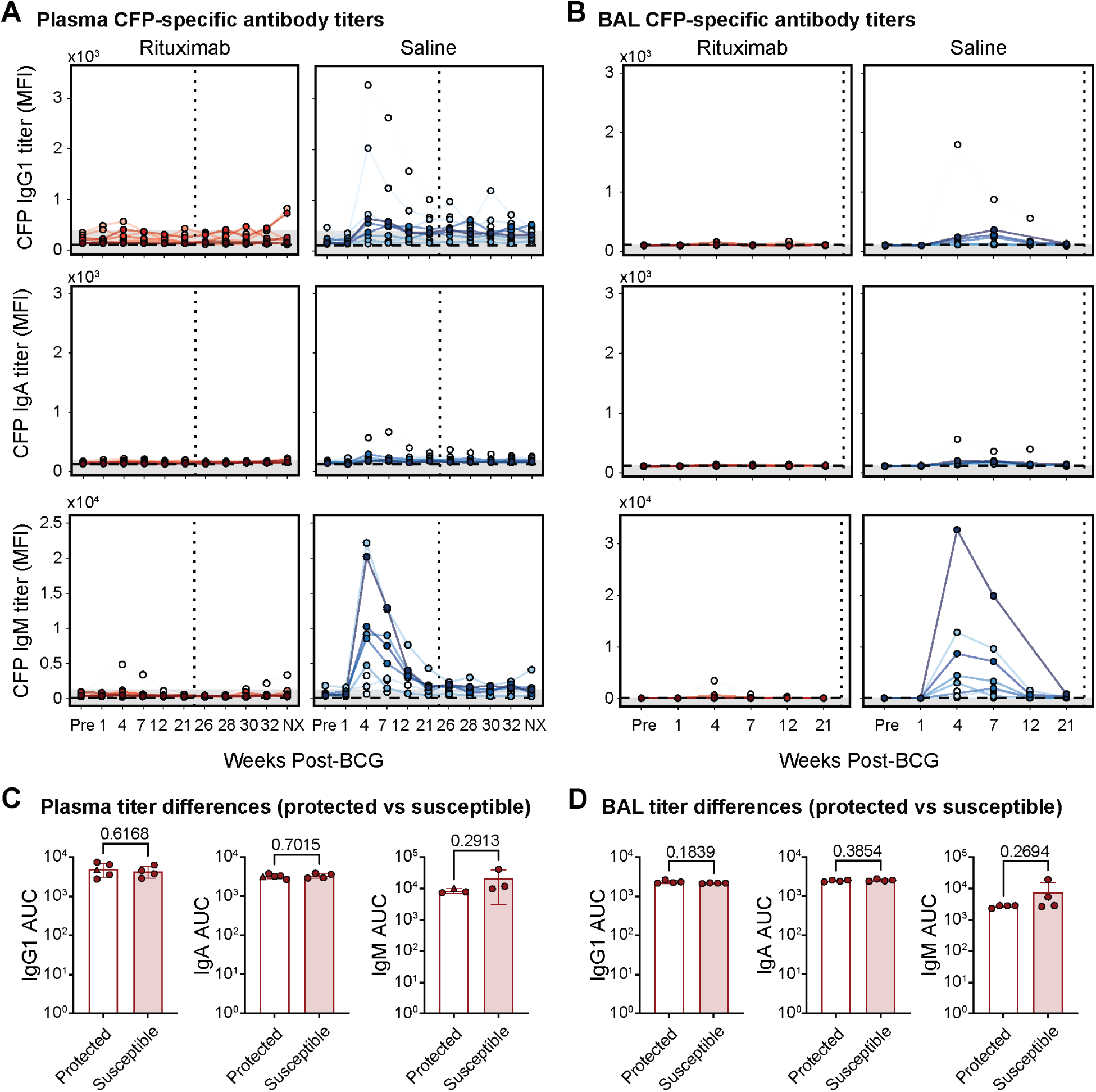
CFP-specific antibody responses. **A**, **B**) IgG1 (top), IgA (middle), and IgM (bottom) antibody titers present in the (**A**) plasma, and (**B**) BAL, of each macaque following IV BCG vaccination specific to CFP measured via Luminex. Y-axis is median fluorescence intensity (MFI). X-axis shows time in weeks, including pre-vaccination at week -4 (Pre) and necropsy at week 38 (NX). Each point represents the duplicate average from a single macaque at the corresponding timepoint. Rituximab-treated macaques are red (left) and saline-treated macaques are blue (right). Vertical dotted line indicates the time of challenge (week 23). Horizontal dashed line is the PBS control. Grey shaded area is the background level set to 2 standard deviations above the mean MFI of the pre-vaccination samples. Measurements at least 2 standard deviations above the mean MFI of the pre-vaccination samples are considered to be above the limit of detection. **C**, **D**) Comparison of CFP-specific antibody titers between protected (total Mtb CFU = 0) and susceptible (total Mtb CFU > 0) Rituximab-treated macaques in the (**C**) plasma, and (**D**) BAL. Area under the curve (AUC) values were calculated using longitudinal Luminex MFI data from the following timepoints: Pre-vaccination (Pre), week 4, 7, 12, and 21. Macaque with 0 Mtb CFU, but residual BCG present in a peripheral lymph node (12620) is represented by the triangle marker. Unadjusted p-values resulting from two-tailed Mann-Whitney U tests are indicated

**Figure S6.**
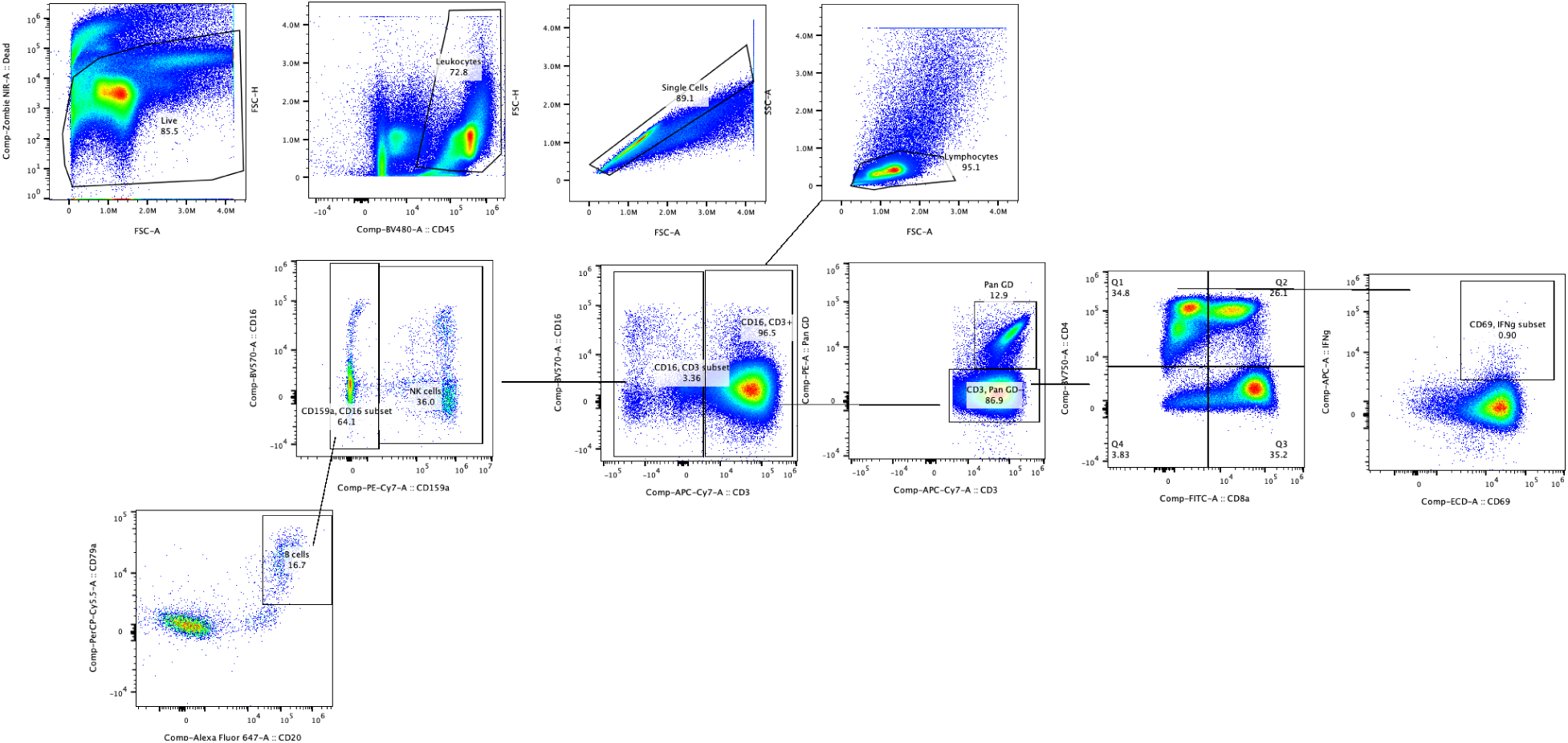
Gating strategy of BAL for flow cytometry.

**Figure S7.**
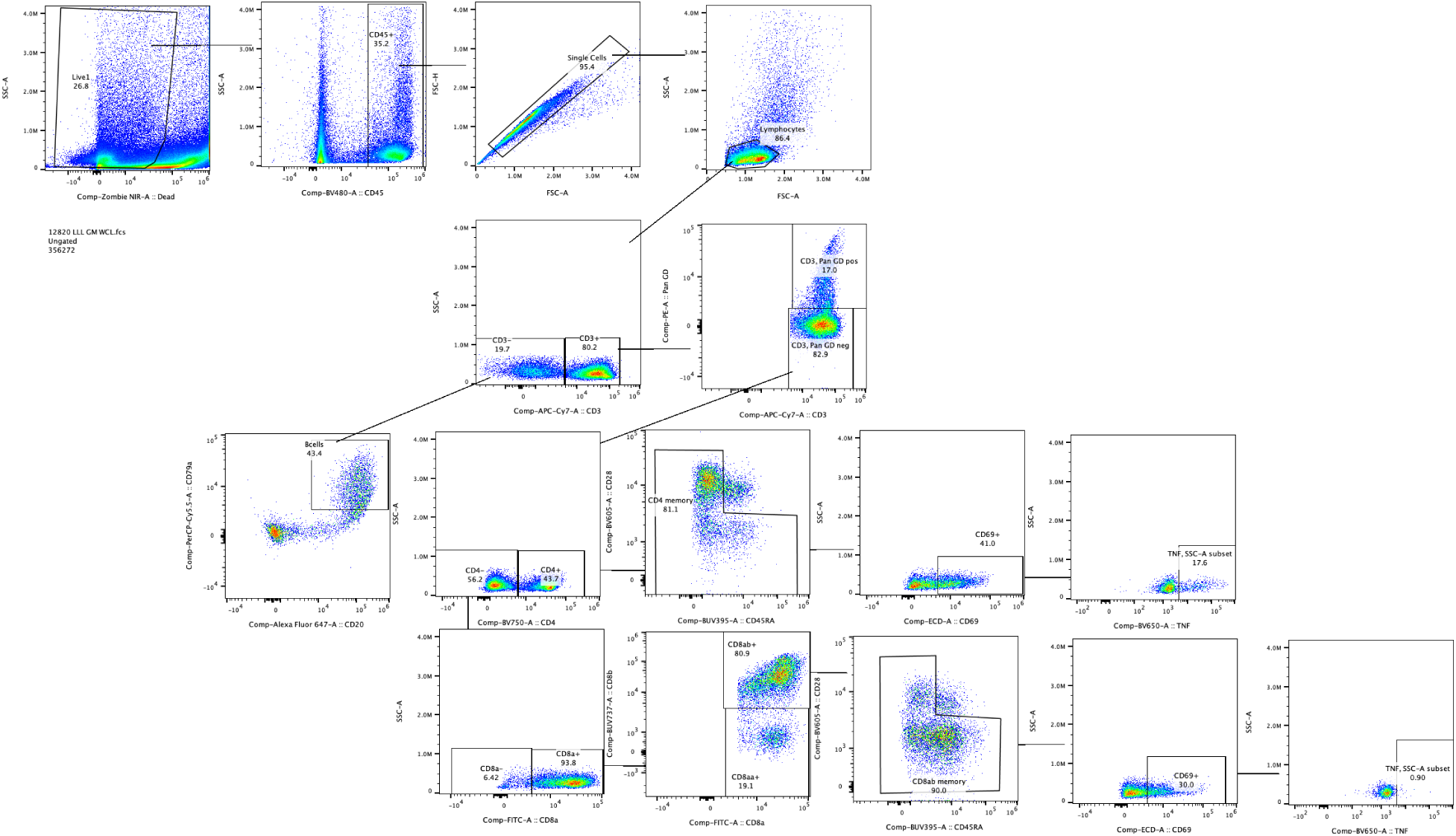
Gating strategy of B cells and memory T cell function in the lung at time of necropsy.

**Table 3.** Flow cytometry antibodies for staining PBMC and Lymph Nodes in Cohorts 1 & 2.

| Clone | Marker | Fluor | Aurora Channel | Vendor | Catalog # | cohort/time point |
| --- | --- | --- | --- | --- | --- | --- |
| 5H9 | CD45RA | BUV395 | UV2 | BD Biosciences | 740315 | cohort 1 week 7-22, all cohort 2 |
| GHI/61 | CD163 | BUV395 | UV2 | BD Biosciences | 745572 | cohort 1 week -4 -4 only |
| ICRF44 | CD11b | BUV563 | UV9 | BD Biosciences | 741357 | cohort 1 and 2 all time points |
| 3.9 | CD11c | BV421 | V1 | BD Biosciences | 565806 | cohort 1 and 2 all time points |
| GB11 | GranzymeB | V450 | V3 | BD Biosciences | 561151 | cohort 1 and 2 all time points |
| 3G8 | CD16 | BV570 | V8 | Biolegend | 302036 | cohort 1 and 2 all time points |
| CD28.2 | CD28 | BV605 | V10 | Biolegend | 302968 | cohort 1 and 2 all time points |
| Ber-ACT8 | CD103 | BV711 | V13 | Biolegend | 350222 | cohort 1 week 7-week 20 |
| 6b11 | iNKT | BV711 | V13 | Biolegend | 342922 | cohort 1 week -4-20 only |
| L200 | CD4 | BV750 | V14 | BD Biosciences | 747202 | cohort 1 and 2 all time points |
| DK25 | CD8a | FITC | B2 | Agilent Technologies | F076501-1 | cohort 1 and 2 all time points |
| HM47 | CD79a | PerCP-Cy5.5 | B9 | ThermoFisher Scientific | 45-0792-42 | cohort 1 and 2 all time points |
| 3D12 | CCR7 | PerCP-eFluor710 | B10 | ThermoFisher Scientific | 46-1979-42 | cohort 1 and 2 all time points |
| 5A6.e9 | Pan gd | PE | YG1 | ThermoFisher Scientific | MHGD04 | cohort 1 and 2 all time points |
| TP1.55.3 | CD69 | ECD | YG3 | Beckman Coulter | 6607110 | cohort 1 and 2 all time points |
| 19.2 | CD206 | PE-Cy5 | YG5 | BD Biosciences | 551136 | cohort 1 and 2 all time points |
| Z199 | CD159a | PE-Cy7 | YG9 | Beckman Coulter | B10246 | cohort 1 and 2 all time points |
| 2H7 | CD20 | AF647 | R2 | Biolegend | 302318 | cohort 1 and 2 all time points |
| H4A3 | CD107a | AF700 | R4 | BD Biosciences | 561340 | cohort 1 week 22, cohort 2 pre BCG |
| LN3 | HLA-DR | AF700 | R4 | ThermoFisher Scientific | 56-9956-42 | cohort 1 week -4-4 only |
| 27E10 | Calprotectin, biotin | AF700 (2 step) | R4 | ThermoFisher Scientific | MA1-40314 | cohort 1 week 7-12 only |
| SP34-2 | CD3 | APC-Cy7 | R7 | BD Biosciences | 557757 (bulk order 624072) | cohort 1 and 2 all time points |
| - | L/D | ZombieNIR | R8 | Biolegend | 423106 | cohort 1 and 2 all time points |

**Table 4.** Flow cytometry antibodies for staining BAL in Cohorts 1 & 2.

| Clone | Marker | Fluor | Aurora Channel | Vendor | Catalog # | cohort/time point |
| --- | --- | --- | --- | --- | --- | --- |
| 5H9 | CD45RA | BUV395 | UV2 | BD Biosciences | 740315 | cohort 1 week 7-22, all cohort 2 |
| GHI/61 | CD163 | BUV395 | UV2 | BD Biosciences | 745572 | cohort 1 week -4 -4 only |
| ICRF44 | CD11b | BUV563 | UV9 | BD Biosciences | 741357 | cohort 1 and 2 all time points |
| 2ST8.5H7 | CD8b | BUV737 | UV14 | BD Biosciences | 748324 | cohort 2 week 12, 22 only |
| 3.9 | CD11c | BV421 | V1 | BD Biosciences | 565806 | cohort 1 and 2 all time points |
| GB11 | GranzymeB | V450 | V3 | BD Biosciences | 561151 | cohort 1 and 2 all time points |
| D058-1283 | CD45 | BV480 | V4 | BD Biosciences | 566145<br>(bulk order<br>(624248) | cohort 1 and 2 all time points |
| BL168 | IL-17 | BV510 | V7 | Biolegend | 512330 | cohort 1 and 2 all time points |
| 3G8 | CD16 | BV570 | V8 | Biolegend | 302036 | cohort 1 and 2 all time points |
| CD28.2 | CD28 | BV605 | V10 | Biolegend | 302968 | cohort 1 and 2 all time points |
| Mab11 | TNF | BV650 | V11 | BD Biosciences | 563418 | cohort 1 and 2 all time points |
| 6b11 | iNKT | BV711 | V13 | Biolegend | 342922 | cohort 1 week -4-20 only |
| BerACT8 | CD103 | BV711 | V13 | Biolegend | 350222 | cohort 1 week 22, cohort 2 pre BCG |
| L200 | CD4 | BV750 | V14 | BD Biosciences | 747202 | cohort 1 and 2 all time points |
| MQ1-17H12 | IL-2 | BV785 | V15 | Biolegend | 500348 | cohort 1 and 2 all time points |
| DK25 | CD8a | FITC | B2 | Agilent Technologies | F076501-1 | cohort 1 and 2 all time points |
| HM47 | CD79a | PerCP-Cy5.5 | B9 | ThermoFisher Scientific | 45-0792-42 | cohort 1 and 2 all time points |
| 3D12 | CCR7 | PerCP-eFluor710 | B10 | ThermoFisher Scientific | 46-1979-42 | cohort 1 and 2 all time points |
| 5A6.e9 | Pan gd | PE | YG1 | ThermoFisher Scientific | MHGD04 | cohort 1 and 2 all time points |
| TP1.55.3 | CD69 | ECD | YG3 | Beckman Coulter | 6607110 | cohort 1 and 2 all time points |
| 19.2 | CD206 | PE-Cy5 | YG5 | BD Biosciences | 551136 | cohort 1 and 2 all time points |
| Z199 | CD159a | PE-Cy7 | YG9 | Beckman Coulter | B10246 | cohort 1 and 2 all time points |
| B27 | IFNg | APC | R1 | Biolegend | 506510 | cohort 1 and 2 all time points |
| 2H7 | CD20 | AF647 | R2 | Biolegend | 302318 | cohort 1 and 2 all time points |
| LN3 | HLA-DR | AF700 | R4 | ThermoFisher Scientific | 56-9956-42 | cohort 1 week -4 -4 only |
| 27E10 | Calprotectin biotin | AF700 (2 step) | R4 | ThermoFisher Scientific | MA1-40314 | cohort 1 week 7-12 only |
| H4A3 | CD107a | AF700 | R4 | BD Biosciences | 561340 | cohort 1 week 22, cohort 2 pre BCG |
| SP34-2 | CD3 | APC-Cy7 | R7 | BD Biosciences | 557757 (bulk order 624072) | cohort 1 and 2 all time points |
| - | L/D | ZombieNIR | R8 | Biolegend | 423106 | cohort 1 and 2 all time points |

**Table 5.** Flow cytometry antibodies for staining lung cells at time of necropsy in Cohorts 1 & 2.

| Clone | Marker | Fluor | Aurora Channel | Vendor | Catalog # | cohort/time point |
| --- | --- | --- | --- | --- | --- | --- |
| 5H9 | CD45RA | BUV395 | UV2 | BD Biosciences | 740315 | cohort 1 and 2 all time points |
| ICRF44 | CD11b | BUV563 | UV9 | BD Biosciences | 741357 | cohort 1 and 2 all time points |
| 2ST8.5H7 | CD8b | BUV737 | UV14 | BD Biosciences | 748324 | cohort 1 and 2<br>all time points |
| 3.9 | CD11c | BV421 | V1 | BD Biosciences | 565806 | cohort 1 and 2<br>all time points |
| GB11 | GranzymeB | V450 | V3 | BD Biosciences | 561151 | cohort 1 and 2<br>all time points |
| D058-1283 | CD45 | BV480 | V4 | BD Biosciences | 566145 (bulk<br>order<br>(624248) | cohort 1 and 2<br>all time points |
| BL168 | IL-17 | BV510 | V7 | Biolegend | 512330 | cohort 1 and 2<br>all time points |
| 3G8 | CD16 | BV570 | V8 | Biolegend | 302036 | cohort 1 and 2<br>all time points |
| CD28.2 | CD28 | BV605 | V10 | Biolegend | 302968 | cohort 1 and 2<br>all time points |
| Mab11 | TNF | BV650 | V11 | BD Biosciences | 563418 | cohort 1 and 2<br>all time points |
| Ber-ACT8 | CD103 | BV711 | V13 | Biolegend | 350222 | cohort 1 and 2<br>all time points |
| L200 | CD4 | BV750 | V14 | BD Biosciences | 747202 | cohort 1 and 2<br>all time points |
| MQ1-17H12 | IL-2 | BV785 | V15 | Biolegend | 500348 | cohort 1 and 2<br>all time points |
| DK25 | CD8a | FITC | B2 | Agilent<br>Technologies | F076501-1 | cohort 1 and 2<br>all time points |
| HM47 | CD79a | PerCP-Cy5.5 | B9 | ThermoFisher<br>Scientific | 45-0792-42 | cohort 1 and 2<br>all time points |
| 3D12 | CCR7 | PerCP-eFluor710 | B10 | ThermoFisher<br>Scientific | 46-1979-42 | cohort 1 and 2<br>all time points |
| 5A6.e9 | Pan gd | PE | YG1 | ThermoFisher<br>Scientific | MHGD04 | cohort 1 and 2<br>all time points |
| TP1.55.3 | CD69 | ECD | YG3 | Beckman Coulter | 6607110 | cohort 1 and 2<br>all time points |
| H4A3 | CD107a | PE-Cy5 | YG5 | BD | 555802 | cohort 1 and 2<br>all time points |
| Z199 | CD159a | PE-Cy7 | YG9 | Beckman Coulter | B10246 | cohort 1 and 2<br>all time points |
| B27 | IFNg | APC | R1 | Biolegend | 506510 | cohort 1 and 2<br>all time points |
| 2H7 | CD20 | AF647 | R2 | Biolegend | 302318 | cohort 1 and 2<br>all time points |
| SP34-2 | CD3 | APC-Cy7 | R7 | BD Biosciences | 557757 (bulk<br>order<br>624072) | cohort 1 and 2<br>all time points |
| - | L/D | ZombieNIR | R8 | Biolegend | 423106 | cohort 1 and 2<br>all time points |

